# ADNKA overcomes SARS-CoV2-mediated NK cell inhibition through non-spike antibodies

**DOI:** 10.1101/2021.04.06.438630

**Authors:** CA Fielding, P Sabberwal, JC Williamson, EJD Greenwood, TWM Crozier, W Zelek, J Seow, C Graham, I Huettner, JD Edgeworth, D Price, BP Morgan, K Ladell, M Eberl, IR Humphreys, B Merrick, K Doores, SJ Wilson, PJ Lehner, ECY Wang, RJ Stanton

## Abstract

The outcome of infection is dependent on the ability of viruses to manipulate the infected cell to evade immunity, and the ability of the immune response to overcome this evasion. Understanding this process is key to understanding pathogenesis, genetic risk factors, and both natural and vaccine-induced immunity. SARS-CoV-2 antagonises the innate interferon response, but whether it manipulates innate cellular immunity is unclear. An unbiased proteomic analysis determined how cell surface protein expression is altered on SARS-CoV-2-infected lung epithelial cells, showing downregulation of activating NK ligands B7-H6, MICA, ULBP2, and Nectin1, with minimal effects on MHC-I. This correlated with a reduction in NK cell activation, identifying a novel mechanism by which SARS-CoV2 antagonises innate immunity. Later in the disease process, strong antibody-dependent NK cell activation (ADNKA) developed. These responses were sustained for at least 6 months in most patients, and led to high levels of pro-inflammatory cytokine production. Depletion of spike-specific antibodies confirmed their dominant role in neutralisation, but these antibodies played only a minor role in ADNKA compared to antibodies to other proteins, including ORF3a, Membrane, and Nucleocapsid. In contrast, ADNKA induced following vaccination was focussed solely on spike, was weaker than ADNKA following natural infection, and was not boosted by the second dose. These insights have important implications for understanding disease progression, vaccine efficacy, and vaccine design.

## Introduction

COVID-19 studies have focused on interferon responses during the early innate phase of infection, and neutralising antibodies and virus-specific T-cells during the adaptive phase. In contrast, and despite their importance in antiviral protection^1^, considerably less is known about the role of Natural Killer (NK) cells in infection. NK cells bridge the innate and adaptive responses, and individuals with NK cell deficiencies suffer severe viral infections^2^. NK cells respond rapidly to viruses, directly killing infected cells by releasing cytotoxic granules. They promote inflammation through the release of TNFα and IFNγ, and influence the induction of both B- and T-cell responses. Once virus-specific antibodies are induced, antibody-dependent NK activation (ADNKA) is generated through antibody linking cell-surface viral antigens with NK cell Fc receptors and subsequent antibody dependent cellular cytotoxicity (ADCC).

The majority of studies on NK cells have focussed on their adaptive roles in established COVID19 infection, with highly activated NK cells seen in both the periphery and the lung^3^. The genetic loss of NKG2C (a hallmark of adaptive NK cells, capable of enhanced ADCC) is also a risk factor for severe disease^4^, implying a protective role in COVID19 infection. In support of this, in animal models, Fc-mediated functions correlate with protection in vaccination^5^, and monoclonal antibody-mediated control of SARS-CoV2 infection is enhanced by Fc-dependent mechanisms^6–11^, which enable control of disease even in the absence of neutralising activity^12^. Furthermore, although neutralising monoclonal antibodies are effective when administered prophylactically, Fc-activity is required for efficacy when administered to animals therapeutically^11^.

These animal studies demonstrate potentially important roles for adaptive NK cell responses in COVID19. Despite this, our understanding of ADCC following infection and vaccination in humans remains limited. Studies to date have assumed spike protein is the dominant ADCC target, and have tested plate-bound protein or transfected cells^13–22^. Using these systems, they have shown that ADNKA-inducing antibodies are generated following SARS-CoV2 infection^14,17–19^, are induced by vaccination in humans^13,16^, and can be mediated by a subset of neutralising monoclonal antibodies targeting the spike protein^20–22^. However, these responses have not been tested against SARS-CoV2 infected cells; the presence, abundance, accessibility, and conformation of viral glycoproteins can be very different during productive infection. Furthermore, although viral entry glycoproteins such as spike can be found on the infected cell surface, and can mediate ADCC, other viral proteins can be the dominant mediators of ADCC during infection; indeed, we have shown that in other viruses, non-structural proteins are often the dominant targets^23^.

In addition, there is little information on the NK cell response to SARS-CoV2 infected cells during the innate phase of infection, despite this process being important to virus pathogenesis; successful viruses must counteract robust NK activation to establish infection, and their ability to do this is critical to their ability to cause disease^24,25^. Whether SARS-CoV2’s ability to infect humans is dependent on viral evasion of innate cellular immunity, is unknown. NK cell activation is complex and depends on the balance of ligands for a wide range of inhibitory or activating receptors^26^. These ligands can be induced on the surface of target cells in response to stress, infection, or transformation. For example, the stress ligands MICA, MICB, and ULBP1-6, all bind to the ubiquitously expressed activating receptor NKG2D. To limit NK cell activation, many viruses manipulate these NK activating ligands, reducing NK-cell mediated control^24^. Whether similar manipulations underlie the ability of SARS-CoV2 to cause disease in humans remains unclear.

To address these critical gaps in our understanding of the SARS-CoV2-specific NK cell response, we used quantitative proteomics to determine how SARS-CoV2 infection affected cell surface protein expression^27^. We found that SARS-CoV2-infected cells downregulate multiple activating NK ligands, resulting in a reduced NK cell response to infected cells. This suggests a novel mechanism by which SARS-CoV2 counteracts innate immunity early in infection, to establish disease. The development of humoral immunity following natural infection resulted in robust ADNKA against infected cells, and high levels of proinflammatory cytokines; these responses persisted for at least 6 months in most patients. Proteomics identified four cell-surface viral proteins capable of driving this response, leading to the surprising discovery that ADNKA is not primarily driven by spike antibody, but was dominated by antibody to other viral proteins such as nucleocapsid, membrane and ORF3a; this defines a novel role for antibodies targeting the immunodominant nucleocapsid protein. In agreement with these findings, and in contrast to studies using plate-bound protein, spike was a poor activator of ADNKA following vaccination when tested using infected cells. This implies that the breadth and magnitude of ADNKA responses is limited in current vaccination strategies. The inclusion of antigens such as nucleocapsid would recruit additional effector mechanisms, and may help maintain vaccine efficacy following mutation of spike in naturally circulating virus variants.

## Results

### SARS-CoV-2 remodels the Plasma Membrane Proteome

Immune cells recognise and interact with infected cells through cell surface ligands. We therefore wanted to gain a comprehensive and unbiased overview of how SARS-CoV-2 infection changes the plasma membrane protein landscape. We performed plasma membrane enrichment through selective aminooxy-biotinylation (Plasma Membrane Profiling; PMP), a methodology that we previously developed^28,29^ and have validated extensively against HIV^30^, HCMV^23,29,31–33^, and HSV^34^ infection.

High infection rates are critical to minimise confounding effects from bystander cells. We therefore applied PMP to SARS-CoV-2 infected A549-ACE2-TMPRSS2 (AAT) cells, which are highly permissive to SARS-CoV-2^27^ and infection at a high multiplicity of infection (MOI) results in over 90% infection by 24h (Fig. 1A). We utilised TMT based quantification to compare PM protein abundance in uninfected cells, and cells infected for 12, 24, and 48 h (Fig 1B). In total, 4953 proteins were quantitated, including 2227 proteins with gene ontology annotation associated with the cell surface and plasma membrane (Fig S1A). Importantly however, the plasma membrane annotated proteins made up nearly 80% of the total protein abundance, indicating a high degree of enrichment for plasma membrane proteins, comparable with our previous PMP datasets^23,28–34^. The detection of low abundance proteins that are not plasma-membrane annotated reflects a combination of incorrect annotations, and contaminating intracellular proteins that were incompletely removed following PMP.

**Fig 1.**
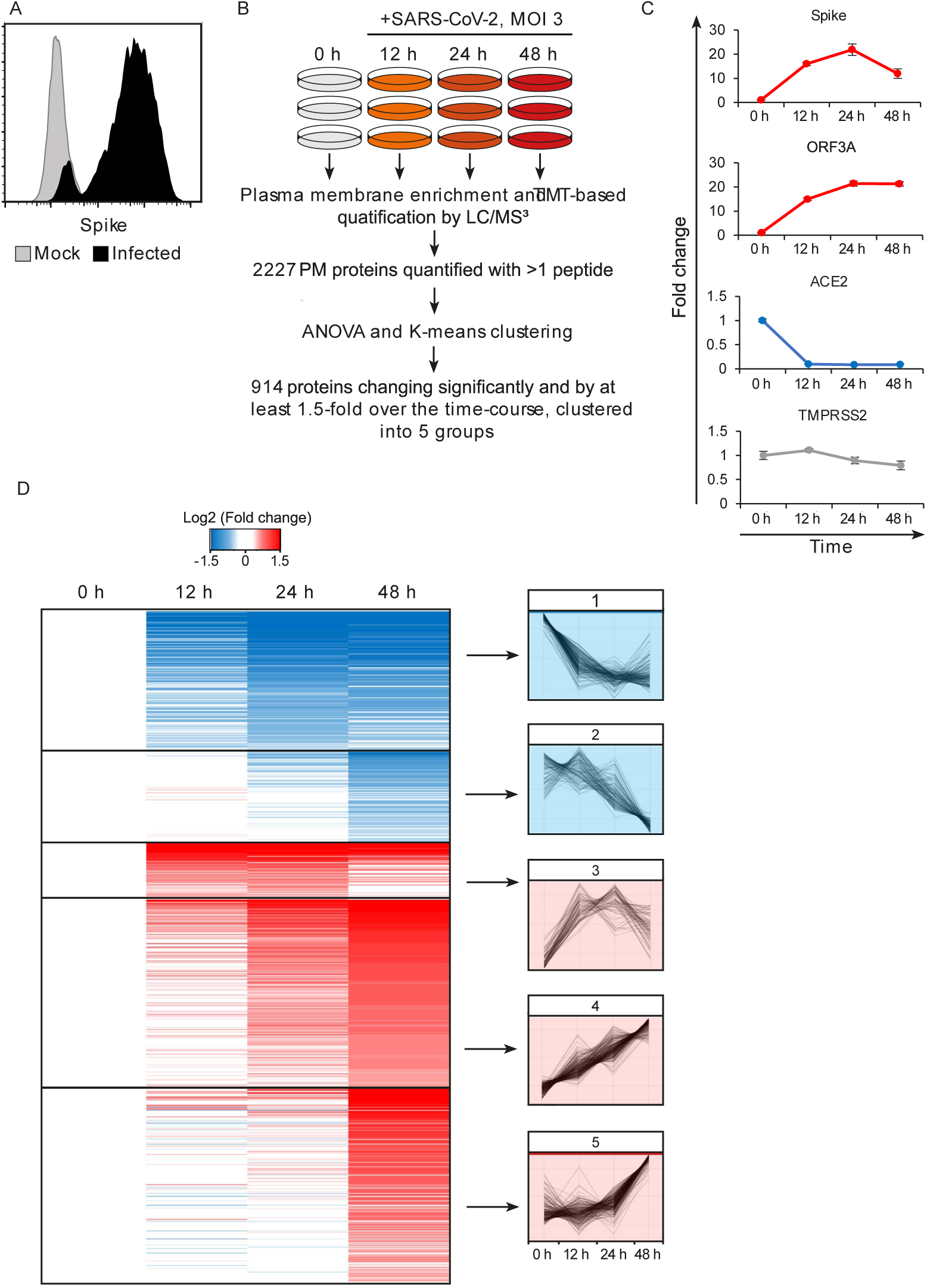
SARS-CoV-2 remodels the Plasma Membrane Proteome. (A) AAT cells were infected with SARS-CoV2 in biological triplicate (MOI=5). 24h later they were detached with trypsin, fixed and permeabilised, stained for Spike protein, and analysed by flow cytometry. (B) Schematic of plasma membrane profiling and analysis pathway. C. Examples of temporal profiles of viral and cellular genes, fold change is compared to 0 h timepoint. Data points show mean ±SD. D. Left – heat map of the 914 significantly changing proteins clustered by k-mean, colour indicates log2 fold change compared to 0 h, right, Z-score normalised temporal profiles of proteins within each cluster.

Five SARS-CoV-2 viral proteins were detected, including spike, ORF3A, and membrane, which have previously been annotated as having a cell surface localisation (Fig. 1C). Consistent with prior reports, the viral receptor ACE2 was strongly downregulated after infection^35^, while TMPRSS2 expression was unaffected (Fig. 1C). Overall, of the 2227 PM proteins quantified with more than one peptide, 914 showed changes which were both significant and with a greater than 1.5-fold over the course of the experiment. Clustering of proteins by their temporal pattern of change has previously shown utility in defining classes of regulated proteins and predicting underlying mechanisms^30,36,37^. Here, we used K-means clustering to define 5 groups of significantly altered proteins by their temporal profile.

To assess the innate immune response triggered by SARS-CoV-2 infection, the clusters were interrogated for the presence of a predefined list of interferon alpha-inducible genes. Thirteen of the 23 detected PM genes associated with the interferon response fell into cluster 5 (Fig S1B), which describes proteins with an increase in expression at 48 h, examples shown in Fig S1C. As the SARS-CoV-2 viral replication cycle is completed in less than 24 h^38^, SARS-CoV-2 is able to evade innate immune recognition and/or supress IFN signalling until long after virus egress has been achieved.

To identify potential innate and adaptive immune evasion strategies employed by SARS-CoV-2, we focused on cluster 1, which describes proteins downregulated from early in the time course. This cluster was subject to gene ontology and pathway analysis to define proteins related by functional and structural similarities (Fig 2). Several classifications enriched in this cluster are particularly relevant, including the downregulation of several cytokine receptors, including IFNAR1. Much of the SARS-CoV-2 genome is dedicated to antagonising a type I interferon response within an infected cell^38^ and the plasma membrane downregulation of IFNAR1 presents a previously undescribed strategy to prevent signalling from exogenous IFN.

**Figure 2.**
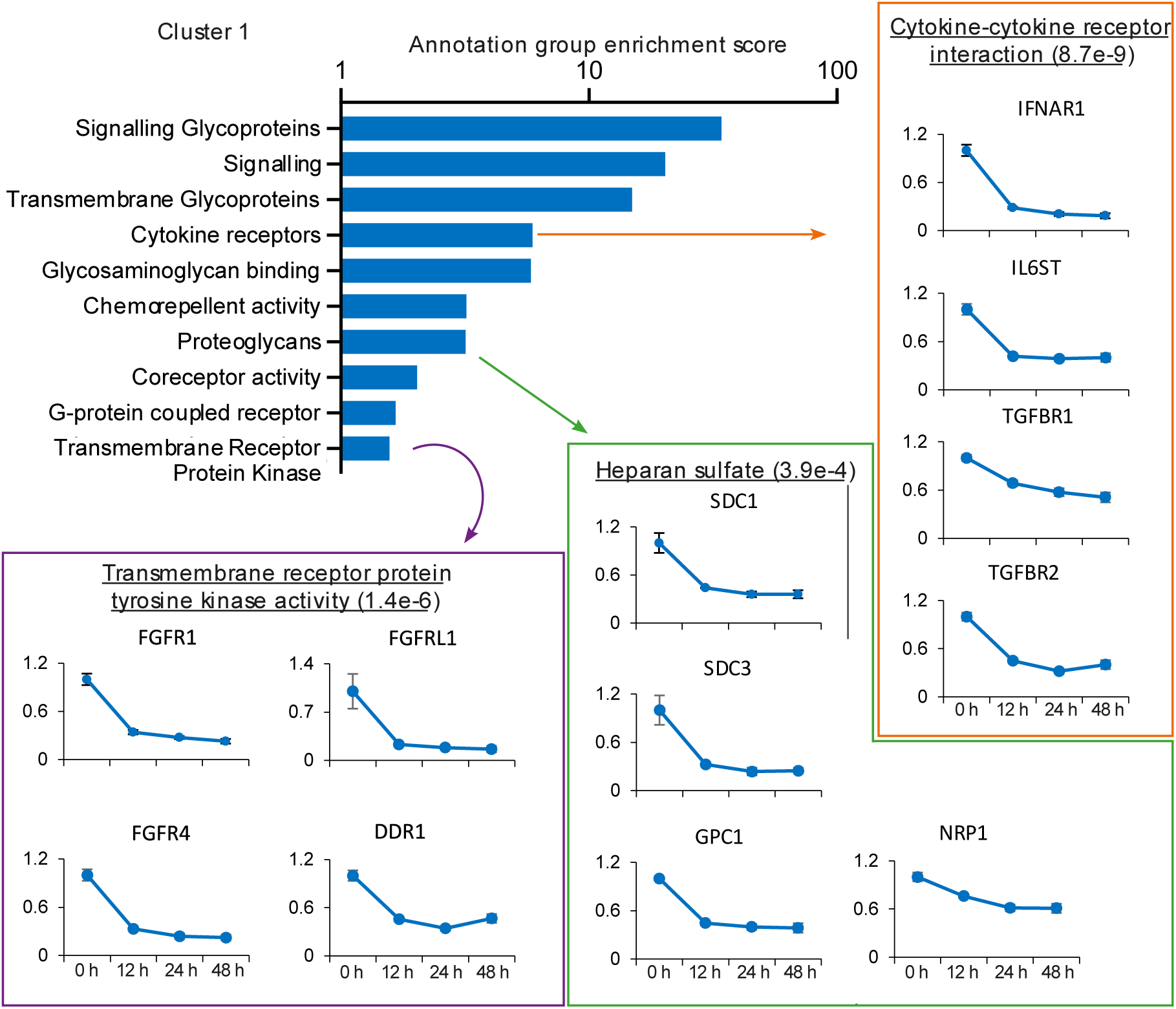
SARS-CoV2 manipulates multiple classes of plasma membrane proteins. (A) Enriched gene ontology and pathway annotations of proteins downregulated in temporal cluster 1 were condensed to groups of related terms, bar chart shows the enrichment score of these groups. Examples of annotation terms falling within each group are shown underlined with Bejamani-Hochberg adjusted P-values in parentheses, alongside four examples of proteins for each annotation.

Also of note is the downregulation of heparan sulfate proteoglycans, the syndecan (SDC), glypican (GPC) and neurophillin (NRP) proteins, which together encompass the families of heparan sulfate modified proteins expressed on epithelial cells^39^. NRP1 is a cell entry factor for SARS-CoV-2^40,41^, and SARS-CoV-2 spike is also reported to bind heparan sulphated proteins^42^. The decrease in cell surface expression of heparan sulfated proteins may therefore enable the egress of viral particles and prevent super-infection, and is reminiscent of the capacity of other viruses to downregulate both primary viral receptors and co-receptors^43^.

### SARS-CoV2 modulates multiple NK cell ligands, and inhibits NK cell activation

Our experience with PMP has emphasised the modulation of NK ligands as an important immune evasion mechanism^31,37^. As these ligands are poorly annotated in public databases, we investigated the PMP dataset for previously described ligands^26^, of which 18 were detected (Fig S2). MHC-I proteins HLA-B to HLA-E were upregulated in cluster 5, likely reflecting a late IFN-mediated upregulation, while HLA-A was unchanged. By contrast, the activating NK-cell ligands MICA, ULBP2, B7-H6 (NCR3LG1), and Nectin1 (which is inhibitory in mice, but recently reported as activating in humans^44^) were significantly downregulated in cluster 1. This was confirmed by flow cytometry (Fig 3A). These responses were dependent on virus replication, because no alterations in ligand abundance were observed when virus was heat-inactivated (Fig S3A).

**Figure 3.**
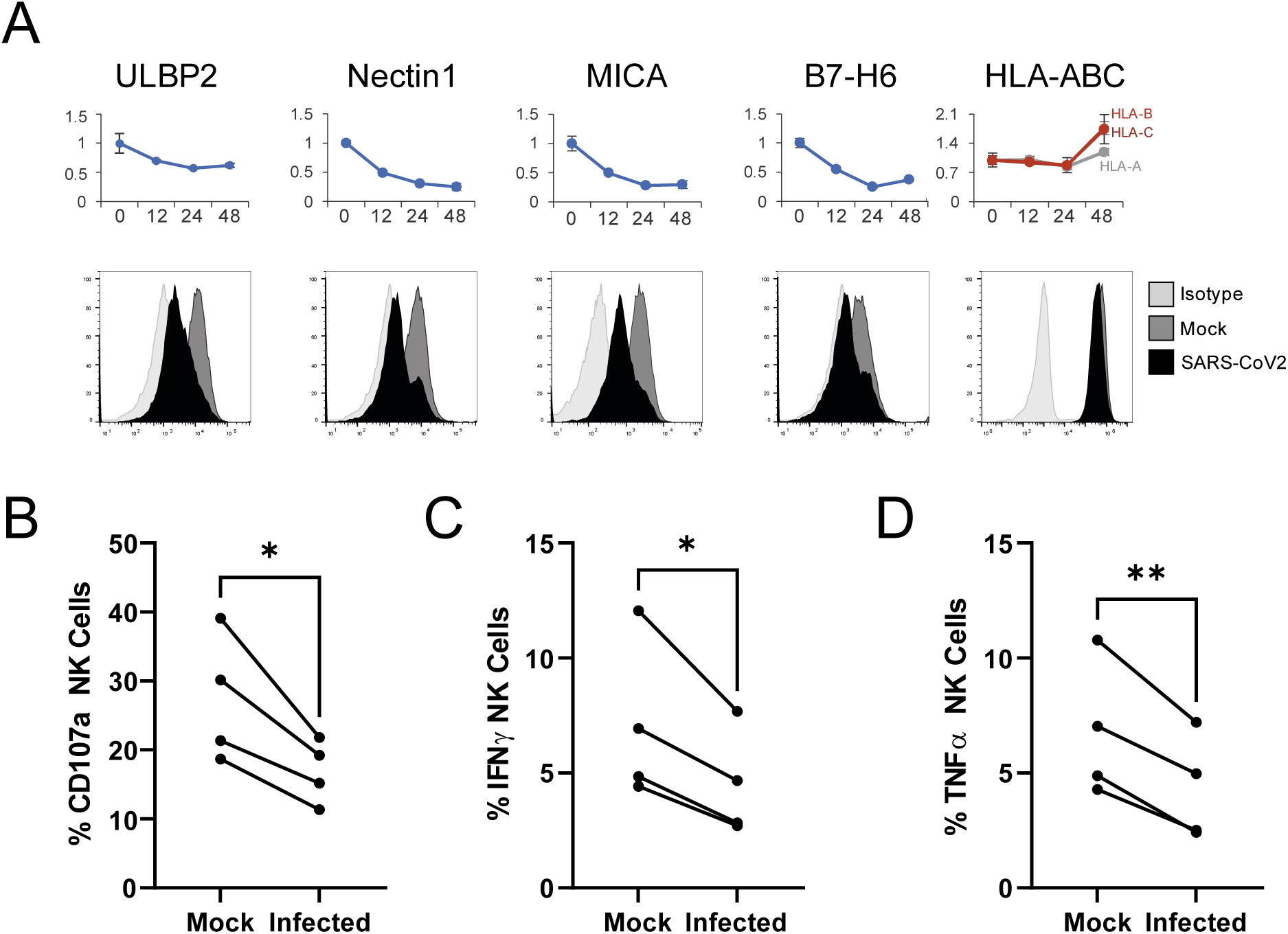
SARS-CoV2 infection leads to downregulation of multiple NK ligands, inhibiting NK activation. (A) AAT cells were either mock infected, or infected with SARS-CoV2 for 24h (MOI=5), detached using TrypLE, then stained for the indicated NK cell receptors before being analysed by flow cytometry (bottom). Plots for the same proteins from PMP are included for reference (top) (B) AAT cells were either mock infected, or infected with SARS-CoV2 for 24h (MOI=5), detached with TrypLE and mixed with interferon stimulated PBMC for 5h in the presence of golgistop, golgiplug, and CD107a antibody, before staining for CD3/CD56 and Live/Dead Aqua. Cells were then fixed, permeabilised, and stained for TNFα and IFNγ. Cells were gated on live CD57+ NK cells, and the percentage of cells positive for CD107a, TNFα and IFNγ calculated. Individual assays were run in technical triplicate, with data shown from four assays using different donors. Kruskal–Wallis, *p<0.05, **p<0.01.

To determine how these changes affected NK cell activation, we incubated SARS-CoV2 infected cells with PBMC, and measured (i) degranulation of CD3^-^CD56^+^ NK cells (Fig 3B), (ii) production of IFNγ (Fig 3C) and (iii) TNFα (Fig 3D, Fig S3B). Consistent with data using other viruses^23^, CD107a was recirculated on a higher proportion of NK cells than IFNγ and TNFα. Nevertheless, all three markers of NK activation were significantly reduced in SARS-CoV2 infected versus mock infected cells, suggesting that the virus has evolved to inhibit NK cell responses. The NK cell recognition system is highly divergent between species^45^, thus the capacity of a virus that is endogenous to bats to manipulate human NK ligands is remarkable. We therefore interrogated the bat genome to determine if it encoded homologues of these ligands. All four have homologues in the horseshoe bat, with one ligand (Nectin1) showing extremely high identity (>96%) to the human gene (Table 1). Thus, control of NK activation may display important similarities between species. It will be crucial to determine whether NK-evasins can function across species barriers, and may therefore play a role in enabling successful zoonotic transfer without the need for extensive adaptation to a new host^46^.

**Table 1.** Homology of NK ligands from humans with those of the horseshoe bat

| Gene | Homo sapiens ID | Rhinolophus ferrumequinum ID | % Identity |
| --- | --- | --- | --- |
| **B7-H6** | XP_011518377 | XP_032975102 | 52.19 |
| **Nectin 1** | NP_002846 | XP_032976402 | 96.12 |
| **MICA** | NP_000238 | XP_032955521 | 43.41 |
| **ULBP2** | NP_079493 | XP_032959100 (ULBP1) | 42.26 |
|  |  | XP_032958641 (ULBP3) | 44.86 |

### NK-cell evasion is overcome by antibody-dependent activation

In addition to their role in innate immunity through interactions with activating and inhibitory NK receptors early in infection, NK cells play additional roles following the development of adaptive immunity via ADNKA/ADCC, in which CD16 interacts with the Fc portion of antibodies bound to targets on the surface of infected cells. To analyse this aspect of NK function, PBMC were incubated with infected cells in the presence of sera from individuals that were seronegative or seropositive for SARS-CoV2. Certain NK cell subsets degranulate more effectively in response to antibody bound targets. These NK cells can be differentiated by the presence of NKG2C and CD57 on the cell surface^47^. However, NKG2C is only found in a subset of donors who are HCMV seropositive, and a proportion of these individuals are NKG2C^null^ due to a deletion in the KLRC2 gene. As a result, NKG2C cannot be used as a marker in all donors. We have previously shown that CD57 is sufficient to identify higher responding NK cells in all donors, and activation of this population correlates with the ability of antibodies to mediate ADCC^48,49^. We therefore focussed on this population here; identical patterns of activation were observed in the CD57 negative population, albeit of slightly smaller magnitude (not shown). Non-specific activation was controlled for by testing against mock-infected cells throughout.

Degranulation (Fig 4A), as well as production of IFNγ (Fig 4B) and TNFα (Fig 4C), were all significantly increased in the presence of serum containing anti-SARS-CoV2 antibodies, across a range of different NK donors (Fig 4D-F). ADNKA was dependent on virus replication, since heat inactivated virus did not lead to serum-dependent NK degranulation (Fig S4A). NK cells were activated equivalently whether purified NK cells or PBMC were used (Fig S4B), thus these responses are due to direct stimulation of NK cells, rather than other cells responding to antibodies and releasing NK stimulating cytokines. Viral immune evasion can therefore be overcome through antibody dependent mechanisms, suggesting that in addition to neutralising virus, SARS-CoV2 antibodies may aid virus clearance through ADCC.

**Figure 4.**
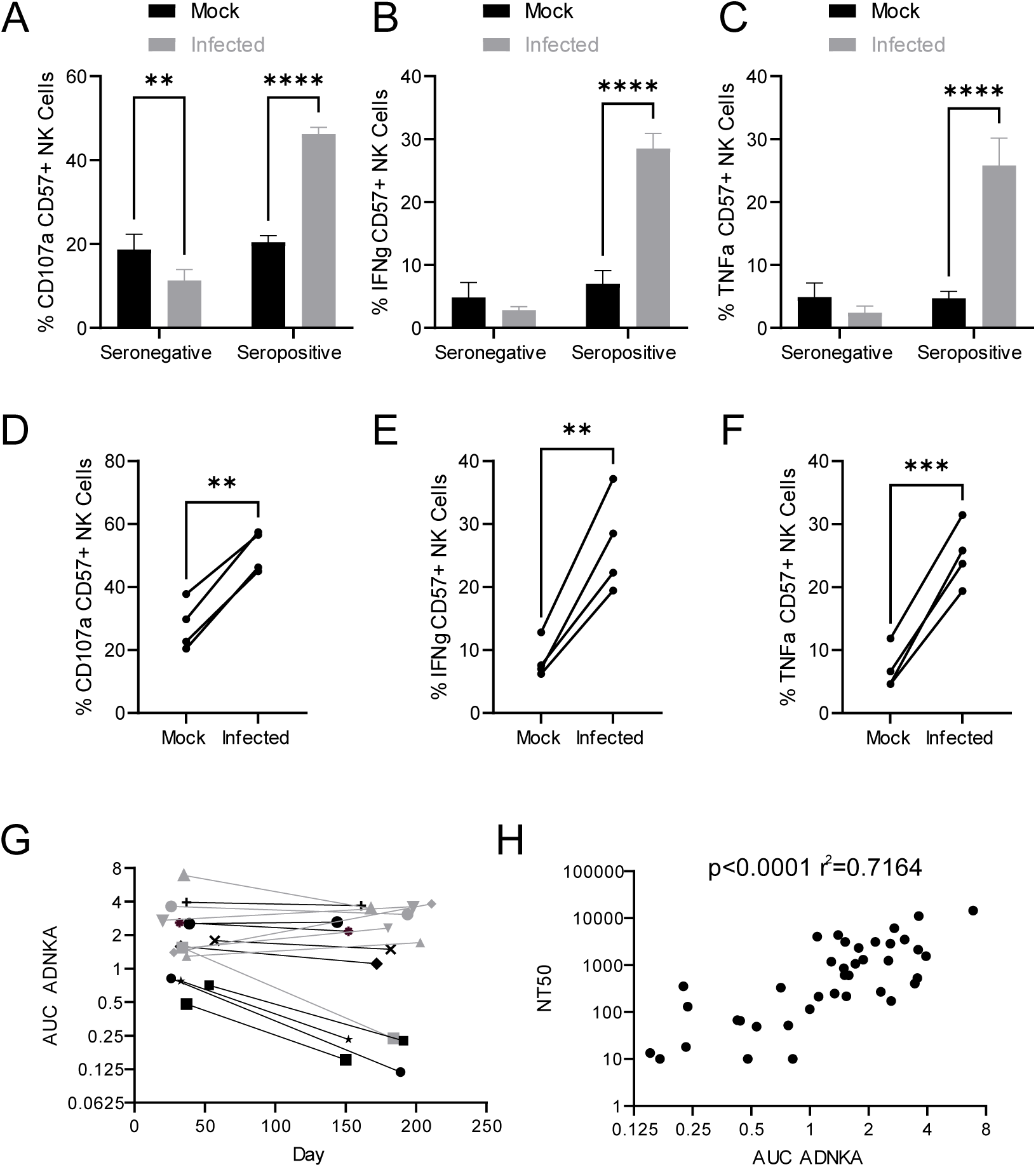
SARS-CoV2 inhibition of NK activation can be overcome via ADNKA. AAT cells were either mock infected, or infected with SARS-CoV2 for 24h (MOI=5), detached using TrypLE, then mixed with PBMC in the presence of golgistop, CD107a antibody, and serum from donors who were seronegative or seropositive for SARS-CoV2. After 5h, cells were stained for CD3, CD56, CD57, and live/dead aqua, then analysed by flow cytometry. (A-F) assays were performed using 1% serum, and the percentage of CD57+ NK cells positive for CD107a (A, D), TNFα (B, E), and IFNγ (C, F) were calculated. Individual donors were performed in technical triplicate (A-C), 2-way ANOVA **p<0.01, ****p<0.0001. Assays from multiple donors were also compared (D-F), Kruskal–Wallis **p<0.01, ***p<0.001. (G-H) assays were performed using a serial 3-fold dilution of serum, starting from 3.3%, then the area under the curve (AUC) calculated. (G) longitudinal serums from donors who experienced mild (black) or severe (grey) were used. (H) In addition to AUC for ADNKA activity, serums were tested for their ability to neutralise SARS-COV2 infection of VeroE6 cells, and the NT50 calculated, then compared to the AUC for ADNKA; Spearman rank correlation analysis is shown.

The magnitude of the neutralising antibody response to SARS-CoV2 correlates with severity of disease; individuals with more severe disease generate higher levels of neutralising antibody^50–52^. Although memory B-cell responses persist, levels of circulating antibody can decrease over time following resolution of disease, such that in some mildly infected people they become undetectable after a few months^50–52^. To determine whether the same was true of ADCC responses, we assessed ADNKA in a cohort of donors with known disease status, for whom longitudinal sera were available (Fig 4G). ADNKA responses were generally of greater magnitude in individuals with more severe disease, with responses remaining detectable during the 150-200 day follow-up. Amongst those with milder disease, strong initial responses were maintained over time, while weaker responses decreased, becoming virtually undetectable by around 200 days. There was a significant correlation between the ability of antibodies to neutralise SARS-CoV2, and to activate ADNKA in response to SARS-CoV2 (Fig 4H), with a R^2^ value of 0.7. Thus, although the antibodies mediating these activities may be induced in a similar manner, the antibodies mediating the two responses may not be identical.

### Multiple SARS-CoV2 ORFs mediate ADCC

There has been a major focus on spike as an activator of ADCC^13–22^. It is certainly a potential target during infection as, despite the fact that virion particles are bud internally rather than from the cell membrane, both our PMP and flow cytometric analysis detected substantial levels of spike protein on the infected cell surface (Fig 5A, 5C, S4C, S4D). However, our studies with other viruses have demonstrated that although viral proteins involved in virion binding and entry are found on the cell surface, and can be bound by antibody, they are not necessarily the strongest mediators of ADCC activity^23^. PMP also detected Membrane (M), Nucleocapsid (N), ORF3a, and ORF1ab on the cell surface. ORF1ab is expressed as a polyprotein, which is proteolytically cleaved into multiple non-structural proteins (NSPs). Analysis of individual peptides representative of the proteolytically processed NSPs did not reveal any specific NSP enrichment. Furthermore, comparing the relative abundance of viral proteins in PMP, ORF1ab was only present at very low levels (1% of all viral proteins, significantly lower than any other SARS-CoV2 protein). Taken together, it is unlikely that any NSPs are surface-exposed. Nevertheless, we performed bioinformatic analysis of all viral proteins for the presence of potential transmembrane domains or signal peptides, revealing five additional candidate cell-surface proteins. We transfected cells with previously validated expression constructs for each of these genes^53^, and screened them for their ability to activate NK cells in an antibody dependent manner. We used sera from patients with severe covid19 disease, that had previously generated the strongest ADNKA signals against infected cells (Fig 5B). As expected, spike primed a substantial ADNKA response. However, in support of our PMP data, expression of ORF3a, Membrane, and Nucleocapsid, were also capable of priming ADNKA in the presence of serum containing SARS-CoV2 antibodies.

**Figure 5.**
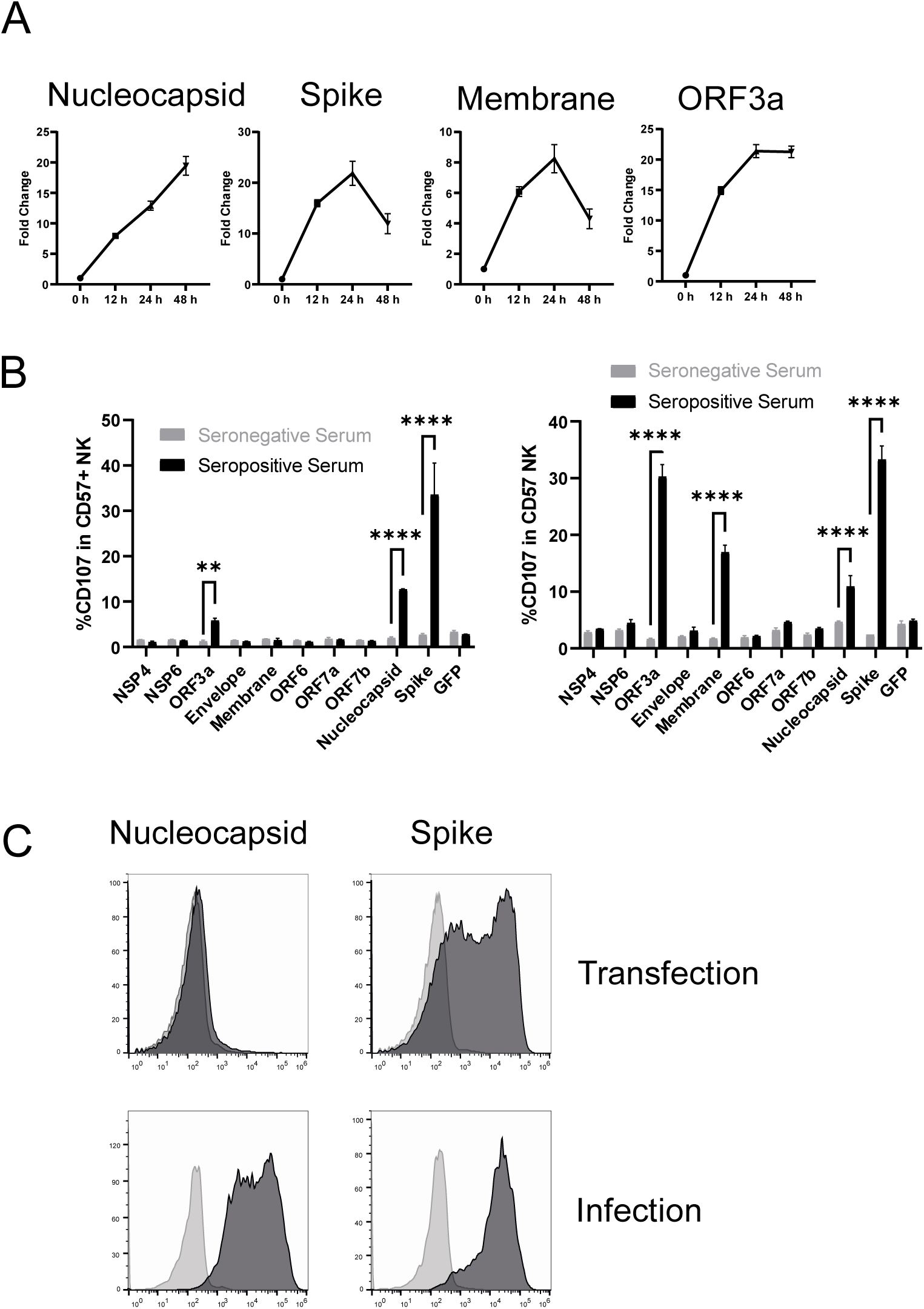
Multiple SARS-CoV2 proteins mediate ADCC. (A) Plots of viral proteins detected in the PMP analysis. (B) 293T cells were transfected with plasmids expressing the indicated SARS-CoV2 proteins for 48h, then mixed with PBMC in the presence of golgistop, CD107a antibody, and 1% serum from donors who were seronegative or seropositive for SARS-CoV2. After 5h, cells were stained for CD3, CD56, CD57, and live/dead aqua, then analysed by flow cytometry. Assays were performed in technical triplicate. Data is shown from two donor serums. **p<0.01, ****p<0.0001, 2-way ANOVA. (C) 293T cells were transfected with plasmid expression SARS-CoV2 nucleocapsid for 48h, or AAT cells were infected with SARS-CoV2 (MOI=5) for 24h, then cells were detached with TrypLE, stained for nucleocapsid or spike, and analysed by flow cytometry.

Interestingly, nucleocapsid readily activated ADNKA despite the extremely low levels of this protein on the surface of transfected cells. We have previously observed the exquisite sensitivity of ADNKA, which can be activated in response to levels of virus protein that are difficult to detect by flow cytometry^23^. We have also shown that antigen levels can be very different in the context of virus infection, compared with transfection^23^. In support of this, nucleocapsid levels were much higher on the surface of infected cells than transfected cells (Fig 5C); detection of nucleocapsid was dependent on virus replication (Fig S4C), and occurred following infection of multiple different cell types (Fig S4D). Given how well even very low levels of surface nucleocapsid activated ADNKA, this suggested that multiple proteins other than spike could be major contributors to ADCC during SARS-CoV2 infection.

### Monoclonal anti-spike antibodies only weakly mediate ADNKA despite binding strongly to infected cells

To determine the relative contribution of spike, and of different antigenic sites on Spike, to ADNKA during SARS-CoV2 infection, we tested a panel of 26 human anti-spike monoclonal antibodies, which had been isolated from naturally infected donors, and cloned as IgG1 constructs^54^. The panel included antibodies targeting the RBD, S1, NTD, and S2 domains, neutralising and non-neutralising, and those targeting multiple distinct epitopes (Table S3).

All 26 antibodies bound efficiently to infected cells, at levels comparable to polyclonal serum (Fig S5). However, only five of these antibodies were capable of triggering degranulation (Fig 6A), of which two triggered TNFα (Fig 6B), and one IFNγ (Fig 6C). Importantly, these NK responses were at significantly lower levels than seen with polyclonal serum from a naturally infected donor. This suggested that, despite high levels of antibody binding, and a substantial ability to activate ADCC when expressed in isolation, spike may be a relatively poor ADCC target in the context of natural infection.

**Figure 6.**
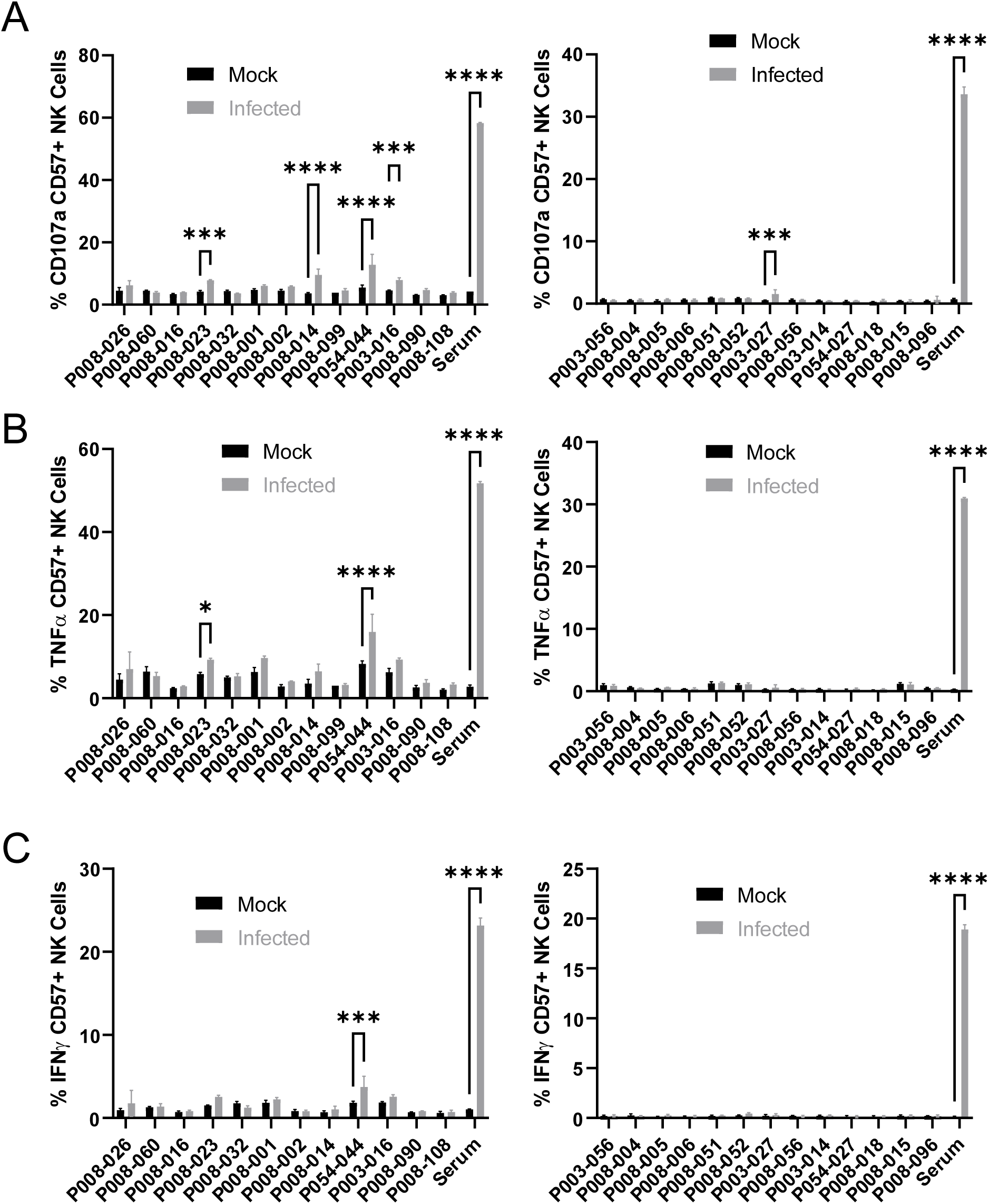
Monoclonal anti-spike antibodies bind infected cells strongly, but only weakly activate ADNKA. AAT cells were either mock infected, or infected with SARS-CoV2 (MOI=5) for 24h, detached with TrypLE and mixed with PBMC and the indicated antibodies, or a serum from a moderate case of covid19. Cells were incubated for 5h in the presence of golgistop, golgiplug, and CD107a antibody, before staining for CD3, CD56, CD57 and Live/Dead Aqua. Cells were then fixed, permeabilised, and stained for TNFα and IFNγ. Cells were gated on live CD57+ NK cells, and the percentage of cells positive for CD107a (A), TNFα (B) and IFNγ (C) calculated. Assays were run in technical triplicate. 2-way ANOVA ***p<0.001, ****p<0.0001.

### Following natural infection, ADNKA responses are dominated by non-spike antibodies

Although our data using monoclonal antibodies indicated that spike was a comparatively weak activator of ADNKA, the antibodies that mediate ADNKA may represent a minor proportion of the overall anti-spike response; we simply may not have captured those antibodies in our panel. To directly address this concern, we specifically depleted anti-spike antibodies from the serum of naturally infected individuals, and assessed changes in activity. ELISA analysis confirmed the effective depletion of spike-specific antibodies (Fig 7A), which was accompanied by a significant loss of neutralisation activity (Fig 7B), across multiple donors (Fig 7C). However, there was no loss of ADNKA activity from any of these samples following spike-antibody depletion (Fig 7D, E), consistent with the results of our monoclonal antibody data. We conclude that spike-specific antibodies dominate the neutralising antibody response, but play a minor roles in the ADCC response following natural infection.

**Figure 7.**
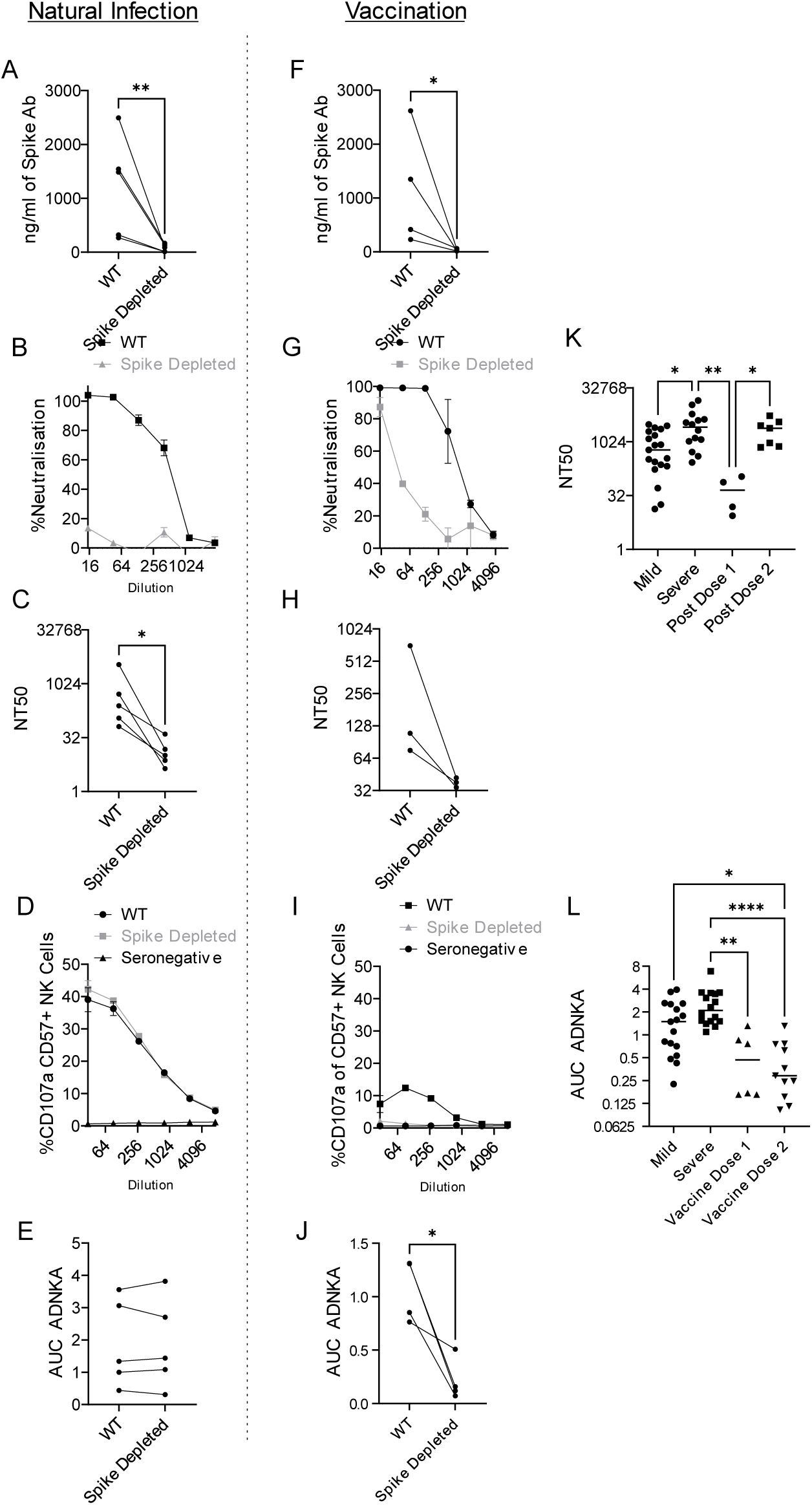
Antibodies targeting spike are weak activators of ADNKA, and the ADNKA response is dominated by non-spike antibodies following natural infection. Serums from individuals naturally infected with SARS-CoV2 (A-E), or donors that had been vaccinated against SARS-CoV2 but were seronegative prior to vaccination (F-J) were depleted of anti-spike antibodies using spike Trimer protein conjugated to magnetic beads. (A, F) ELISA for spike trimer was used to measure levels of antibodies before and after depletion. (B, C, G, H) the ability of the original, or anti-spike depleted, serums to neutralise the ability of SARS-CoV2 to infect VeroE6 cells was determined across a range of concentrations, then NT50 values calculated. Example plots (B, G) and NT50 values for multiple donors (C, H) are shown. (D, E, I, J) AAT cells were either mock infected, or infected with SARS-CoV2 for 24h (MOI=5), detached using TrypLE, then mixed with PBMC in the presence of golgistop, CD107a antibody, and serial dilutions of serum. After 5h, cells were stained for CD3, CD56, CD57, and live/dead aqua, then analysed by flow cytometry for the percentage of CD107a positive CD57+ NK cells. AUC values were then calculated. All serums were additionally tested against mock infected cells to ensure no non-specific NK activation occurred (not shown). Example plots (D, I) and AUC values for multiple donors (E, J) are shown. NT50 (K) and AUC (L) values are also given for samples categorised according to disease or vaccination status. Kruskal–Wallis *p<0.05, **p<0.001.

### Spike-specific antibodies following vaccination are weak mediators of ADNKA

Since non-spike antigens must prime ADCC following natural infection, we investigated individuals who have been vaccinated against spike, but not exposed to the virus - since they will only have spike-specific antibodies. As with serum from naturally infected individuals, ELISA demonstrated effective depletion of spike antibodies (Fig 7F), which was accompanied by a marked loss of virus neutralisation activity (Fig 7G, H). In functional NK assays, these sera were capable of mediating relatively weak ADNKA (Fig 7I, J) and, unlike sera from naturally infected individuals, ADNKA activity was abrogated following depletion of spike antibodies. When we compared responses following natural infection or vaccination, vaccine mediated ADNKA was significantly weaker than responses seen following natural infection, and was not boosted following the second dose of the vaccine (Fig 7K). This was in stark contrast with the neutralisation activity of those same serum samples, which were dramatically increased following the second dose, with activity comparable to patients with severe covid-19 disease (Fig 7L).

Thus, anti-spike antibodies can mediate ADNKA, but this activity appears weaker than responses seen during natural infection, which targets a broader repertoire of viral antigens.

## Discussion

The host innate immune system plays a critical role in both early SARS-CoV2 infection, and later COVID-19 disease. This is emphasised by the identification of multiple interferon antagonists encoded by SARS-CoV2, and the increased disease severity in patients with genetic and acquired interferon deficiencies^38,55–57^. To determine whether the virus evades other components of the host immune system we took a proteomic approach to systematically analyse how SARS-CoV2 infection manipulates the cell surface during the time course of SARS-CoV2 infection. We find that SARS-CoV-2 remodelling of the plasma membrane leads to the downregulation of multiple cell surface immune ligands involved with the interferon response, cytokine function, and NK activation. This may enable SARS-CoV2 to evade numerous effector arms of the host immune system. Of particular interest, downregulation of NK activating ligands inhibits NK cell activation, and is therefore likely to play an important role in the ability of the virus to establish initial infection. Later in disease, following the production of antibody, ADNKA leads to significant degranulation, and production of pro-inflammatory cytokines. Somewhat surprisingly, spike antibody is not responsible for the robust ADNKA which develops after natural infection, and only weak ADNKA is generated following spike vaccination.

Genetic variation in both viruses and their hosts affects the ability of viruses to modulate NK responses, and influence pathogenesis^24,58^. SARS-CoV2 appears to have evolved to antagonise NK activation by targeting the expression levels of multiple activating NK ligands. Nectin-1 is a ligand for CD96/TACTILE^59^, which activates human NK cells^44^, B7-H6 is a ligand for NKp30, while ULBP2 and MICA are ligands for NKG2D. NKG2D is ubiquitously expressed on γδ cells and CD8+, as well as some CD4+, T-cells in addition to NK cells^60^. Targeting NKG2D ligands may therefore enable the virus to impact additional effector mechanisms beyond just NK cells. Amongst NKG2D ligands, MICA is particularly polymorphic (100+ alleles), A549 cells express MICA*001/004^61^, while the truncated MICA*008 is the most common allele worldwide. This diversity is largely driven by co-evolution of humans with virus infections, and reflects the broad repertoire of viral immune-evasins^60^. MICA variation therefore has the potential to play a role in the extremely variable outcome of SARS-CoV-2 infection.

In contrast to the effect on NK ligands, the virus did not significantly downregulate MHC-I, implying that it cannot prevent peptide presentation to antagonise adaptive cytotoxic cellular immunity. Nor do infected cells bind human IgG from seronegative donors, indicating that it does not encode Fc receptors as ADNKA decoys. These observations are consistent with SARS-CoV2 acting as an acute ‘hit and run’ virus, which has replicated and transmitted before host adaptive immunity develops. It will be interesting to determine whether coronaviruses evade innate and adaptive immunity in their original bat host, and whether this contributes to their persistence in that species.

In animal models of vaccination^5^ and monoclonal antibody administration^6–9,11^ against SARS-CoV2, antibody-mediated activation of cellular immunity is a critical component of the protective immune response. However, inflammation dominates severe COVID-19 disease^62,63^. The high levels of TNFα and IFNγ observed following ADNKA are likely to exacerbate disease under these circumstances. In support of this, TNFα and IFNγ correlate with severe disease^64^, and critically ill patients have a higher proportion of afucosylated antibodies that may promote a stronger ADCC response^65^. Given that ADCC can potentially contribute to both protection and immunopathology, it is important to understand what determines the magnitude and quality of the ADCC response. Viral entry glycoproteins are a common focus as mediators of ADCC. E.g. ADCC mediated by Env antibodies have been a major focus in HIV^66^, and gB-specific antibodies have been studied in HCMV^67^. Although antibodies targeting these antigens can clearly bind infected cells, using similar approaches to the present study we recently showed that non-structural accessory proteins were more potent ADCC targets during HCMV infection^23^. This conclusion is reinforced here, where the neutralising antibody response is dominated by spike, but spike antibodies are poor activators of ADNKA, with other viral antigens driving more efficient ADNKA following natural infection. In support of this data, monoclonal anti-spike antibodies were only effective at activating Fc-mediated control of infection in animal models following engineering to enhance activity^12^. Importantly, the ability of monoclonal spike antibodies to bind to infected cells did not correlate with their ability to mediate ADNKA, implying that the epitope, position, or angle, at which the antibody binds to the cell surface is critical; it’s notable that 4/5 anti-spike antibodies that activated ADNKA bound the NTD, suggesting that this site is structurally advantageous for Fc-mediated responses. A similar phenomenon may underly the greater ability of non-spike antigens to generate a ADNKA response. Nucleocapsid, ORF3a, and membrane are all present on the infected cell surface, and can promote ADNKA if appropriate antibodies are present. Antibodies targeting ORF3a and membrane have been reported as present in only a subset of patients, while almost everyone infected with SARS-CoV2 develops antibodies targeting^68^. Nucleocapsid is therefore likely to be the major target for ADCC during natural SARS-CoV2 infection in most people, identifying a potential functional role for antibodies binding this immunodominant target for the first time. Influenza nucleoprotein is also detected at the plasma membrane^69–71^, and antibodies targeting nucleoprotein can mediate ADCC^72–74^. Structural proteins involved in genome packaging may therefore represent a common target for ADCC across multiple virus families.

The identification of the antigens that drive ADCC is important to our understanding of the mechanisms that underlie pathogenesis and disease. A skewing of the spike:nucleocapsid antibody ratio is associated with severe disease^75–77^, where ADCC-mediating nucleocapsid antibodies might be promoting inflammation. Both the breadth of the nucleocapsid antibody response, and the particular epitopes targeted, have also been correlated with disease severity^68,78^. These correlations now need to be followed up functionally, to determine how the epitopes bound by anti-nucleocapsid antibodies affect ADCC. Critically, the same surface-bound antibodies that activate NK cells are likely to promote inflammation by activating myeloid cells and complement^79–81^, both of which are hallmarks of severe Covid19.

Although Fc-dependent mechanisms of cellular immune activation could contribute to immunopathology via multiple mechanisms in severe disease, all patients with mild/asymptomatic disease also developed strong and persistent ADNKA responses. Furthermore, lineage defining mutations in the viral variants of concern B.1.1.7, B.1.351, and P.1, all include non-synonymous mutations in nucleocapsid, which is one of the most variable genes amongst circulating strains^82^, and these nucleocapsid mutations can result in reduced antibody binding^83^. Virus variants may therefore evolve to evade ADCC responses as well as neutralising responses, supporting an important role for ADCC in protection. It is therefore significant that spike-vaccine generated antibody responses were poor ADNKA inducers when tested against infected cells. The addition of other viral proteins such as nucleocapsid to vaccines would engage a wider range of immune effector pathways. This approach might improve efficacy against both viral transmission and disease, resulting in vaccines that are more resistant to viral variants containing mutations that diminish antibody neutralisation^84^.

## Acknowledgements

This work was partly funded by the MRC/NIHR through the UK Coronavirus Immunology Consortium (CiC; MR/V028448/1). RS was supported by the MRC (MR/S00971X/1), Wellcome Trust (204870/Z/16/Z), and Ser Cymru. ECYW was funded by the MRC (MR/P001602/1, MR/V000489/1). PJL was funded by the Wellcome Trust through a Principal Research Fellowship (210688/Z/18/Z), the MRC (MR/V011561/1), the Addenbrooke’s Charitable Trust and the NIHR Cambridge Biomedical Research Centre. KLD was supported by King’s Together Rapid COVID-19 Call, Huo Family Foundation Award, and a Wellcome Trust Multi-User Equipment Grant 208354/Z/17/Z. CG was supported by the MRC-KCL Doctoral Training Partnership in Biomedical Sciences (MR/N013700/1). BM was supported by an NIHR Academic Clinical Fellowship in Combined Infection Training

## Materials and methods

### Cells and viruses

A549 were transduced with lentiviruses expressing human ACE2, and TMPRSS2 (AAT cells), as previously described^27^. The England2 strain of SARS-CoV2 was obtained from Public Health England (PHE), and grown on VeroE6 cells. Virions were concentrated by pelleting through a 30% sucrose cushion to remove contaminating soluble proteins, and titrated by plaque assay on both VeroE6, AAT, and Caco2, as previously described^27^. All cell lines were grown in DMEM containing 10% FCS (Gibco), at 37°C and in 5% CO_2_. Throughout the study, multiple batches of virus were used, and no overt differences were noted between them. For assays, cells were plated out the day before, then infected at a multiplicity of infection (MOI) of 5, for 1h on a rocking platform. The inoculum was removed, and fresh DMEM containing 2% FCS was added, then cells were incubated until the assay. To confirm that phenotypes were dependent on active virus replication, virus was inactivated by heating to 56°C for 30 minutes.

### Plasma membrane profiling

Cell surface proteins were labelled essentially as described^29,85^. Briefly, cells were incubated in a solution containing sodium periodate, aniline and aminooxy biotin to label predominantly sialic acid containing glycans at the cell surface. Cell lysates were then enriched for labelled proteins using streptavidin-agarose beads. After extensive washing trypsin was added to liberate peptides of the enriched proteins. The resulting peptide pools were dried prior to labelling with TMT reagents.

### TMT Labelling and clean-up

Samples were resuspended in 21µL 100mM TEAB pH 8.5. After allowing to come to room temperature, 0.2mg TMT reagents (Thermo Fisher) were resuspended in 9µL anhydrous ACN which was added to the respective samples and incubated at room temperature for 1h. A 3µL aliquot of each sample was taken and pooled to check TMT labelling efficiency and equality of loading by LC-MS. After checking each sample was at least 98% TMT labelled total reporter ion intensities were used to normalise the pooling of the remaining samples such that the final pool was as close to a 1:1 ratio of total peptide content between samples as possible. This final pool was then dried in a vacuum centrifuge to evaporate the majority of ACN form labelling. The sample was acidified to a final 0.1% Trifluoracetic Acid (TFA) (∼200µL volume) and FA was added until the SDC visibly precipitated. 4 volumes of ethyl acetate were then added and the sample vortexed vigorously for 10s. Sample was then centrifuged at 15’000g for 5mins at RT to effect phase separation. A gel loading pipette tip was used to withdraw the lower (aqueous) phase to a fresh low adhesion microfuge tube. The sample was then partially dried in a vacuum centrifuge and brought up to a final volume of 1mL with 0.1% TFA. FA was added until the pH was <2, confirmed by spotting onto pH paper. The sample was then cleaned up by SPE using a 50mg tC18 SepPak cartridge (Waters). The cartridge was wetted with 1mL 100% Methanol followed by 1mL ACN, equilibrated with 1mL 0.1% TFA and the sample loaded slowly. The sample was passed twice over the cartridge. The cartridge was washed 3x with 1mL 0.1% TFA before eluting sequentially with 250µL 40% ACN, 70% ACN and 80% ACN and dried in a vacuum centrifuge.

### Basic pH reversed phase fractionation

TMT labelled samples were resuspended in 40µL 200mM Ammonium formate pH10 and transferred to a glass HPLC vial. BpH-RP fractionation was conducted on an Ultimate 3000 UHPLC system (Thermo Scientific) equipped with a 2.1 mm × 15 cm, 1.7µ Kinetex EVO column (Phenomenex). Solvent A was 3% ACN, Solvent B was 100% ACN, solvent C was 200 mM ammonium formate (pH 10). Throughout the analysis solvent C was kept at a constant 10%. The flow rate was 500 µL/min and UV was monitored at 280 nm. Samples were loaded in 90% A for 10 min before a gradient elution of 0–10% B over 10 min (curve 3), 10-34% B over 21 min (curve 5), 34-50% B over 5 mins (curve 5) followed by a 10 min wash with 90% B. 15s (100µL) fractions were collected throughout the run. Fractions containing peptide (as determined by A280) were recombined across the gradient to preserve orthogonality with on-line low pH RP separation. For example, fractions 1, 25, 49, 73, 97 are combined and dried in a vacuum centrifuge and stored at -20°C until LC-MS analysis. 24 Fractions were generated in this manner.

### Mass Spectrometry

Samples were analysed on an Orbitrap Fusion instrument on-line with an Ultimate 3000 RSLC nano UHPLC system (Thermo Fisher). Samples were resuspended in 10µL 5% DMSO/1% TFA and all sample was injected. Trapping solvent was 0.1% TFA, analytical solvent A was 0.1% FA, solvent B was ACN with 0.1% FA. Samples were loaded onto a trapping column (300µm x 5mm PepMap cartridge trap (Thermo Fisher)) at 10µL/min for 5 minutes at 60 degrees. Samples were then separated on a 75cm x 75µm i.d. 2µm particle size PepMap C18 column (Thermo Fisher) at 55 degrees. The gradient was 3-10% B over 10mins, 10-35% B over 155 minutes, 35-45% B over 9 minutes followed by a wash at 95% B for 5minutes and requilibration at 3% B. Eluted peptides were introduced by electrospray to the MS by applying 2.1kV to a stainless-steel emitter (5cm x 30µm (PepSep)). During the gradient elution, mass spectra were acquired with the parameters detailed in Fig S6 using Tune v3.3 and Xcalibur v4.3 (Thermo Fisher).

### Data Processing

Data were processed with PeaksX+, v10.5 (Bioinformatic Solutions). Processing parameters are shown in detail in Supplementary Fig S7. Briefly, .raw files were searched iteratively in three rounds, with unmatched *de novo* spectra (at 1% PSM FDR) from the previous search used as the input for the next. The three iterations were as follows 1) Swissprot Human (27/03/2020) + common contaminants 2) The same databases as search 1 but permitting semi-specific cleavage 3) trEMBL Human (27/03/2020), with specific cleavage rules. Proteins were then quantified using the parameters outlined in Fig S7. Identified proteins and their abundances were output to .csv format and further subjected to statistical analysis. The mass spectrometry proteomics data have been deposited to the ProteomeXchange Consortium via the PRIDE partner repository^86^ with the dataset identifier PXD025000 and 10.6019/PXD025000. Table S1 contains the analysed data.

### Statistical analysis and K-means clustering

Prior to statistical analysis, proteins were filtered for those quantified across all TMT reporter channels and with more than 1 unique. Only proteins with a plasma membrane related GO:CC annotation as previously described^28^ were then carried forward for statistical analysis. All SARS-CoV-2 proteins quantified with more than one unique peptide were also included. Statistical tests were performed with the aov and p.adjust functions in the R base stats package (version 4.0.3) to calculate Benjamini-Hochberg corrected p-values for changes in protein abundance across the measured time-points.

Proteins with a p-value < 0.05 and a maximum fold-change across the time-course of > 1.5-fold were selected for k-means clustering. Mean intensities of each time-point were scaled using the scale function in base R and utilised to cluster proteins into 5 groups using the kmeans function in R base stats package.

### Functional Annotation Clustering

Assessment of enriched gene annotation terms in temporal cluster one was carried out using the Functional Annotation Clustering tool at DAVID (david.ncifcrf.gov) v6.8^87^, using the default clustering settings for medium stringency and the following libraries: Uniprot UP_Keyword, GOTERM:MF_ALL, BIOCARTA,, KEGG_PATHWAY and REACTOME_PATHWAY. Output clusters were curated with representative names based on the enriched terms within. The background for enrichment was a list of proteins detected in the PMP dataset with more than one peptides and a plasma membrane associated GO:CC term. Functional annotation clusters with an enrichment score of >1.5 are shown, and any clusters in which no individual annotation term had a Benjamini-Hochberg correct p-value of <0.05 were excluded.

### Flow cytometry

Cells were dissociated using TrypLE, then stained with primary antibody for 30 min at 4°C. Following washing, they were incubated with secondary antibody, again for 30 min at 4°C. Cells were washed, fixed in 4% paraformaldehyde, and data collected on an Attune cytometer (Thermofisher). In one experiment (Fig 1A), cells were fixed and permeabilised (Cytofix/Cytoperm, BD) before staining. Antibodies used were against HLA-ABC (W632; AbD Serotec), NCR3LG1/B7-H6 (MAB7144, Biotechne R&D Systems), Nectin 1 (R1.302; Biolegend), MICA (AMO1-100; BAMOMAB), MICB (BMO2-100; BAMOMAB), ULBP2 (BUMO1; BAMOMAB), Spike (1A9; Insight), Nucleocapsid (1C7; Stratech), anti-mouse IgG AF647 (Thermofisher).

### ELISA

To measure levels of Spike antibodies, an ELISA using the Spike Trimer (Acro Biosystems) was used according to manufacturers instructions. Each sample was measured in duplicate, and compared to a standard curve. A pre-pandemic serum was included in each assay to define the cutoff.

### Plasmids and transfections

Lentivirus plasmids encoding each SARS-CoV2 ORF individually, with a C-terminal twin-strep tag, were obtained from Addgene, and have been validated for expression previously^53^. Plasmids were midiprepped (Nucleobond Xtra Midi; Machery-Nagel), and transfected into 293T cells using GeneJuice (Merck) according to manufacturers’ instructions.

### NK activation assays

PBMCs from healthy donors were thawed from liquid N2 storage, rested overnight in RPMI supplemented with 10% FCS, and L-glutamine (2 mM). For assays investigating viral inhibition of NK activation, PBMC were stimulated overnight with IFN-α (1,000 U/ml) to provide a baseline level of activation against which inhibition could be seen. For ADNKA this stimulation is not required, and so cells were used unstimulated^23^. To confirm that phenotypes were due to direct effects on NK cells, NK cells were purified by depletion, using the human NK cell purification kit (Miltenyi) according to manufacturer’s instructions. Target cells were harvested using TrypLE Express (Gibco), preincubated for 30 min with the relevant antibody or serum preparations, then mixed with effectors at an effector:target (E:T) ratio of 10:1 (PBMC) or 1:1 (purified NK cells) in the presence of GolgiStop (0.7 μl/ml, BD Biosciences), Brefeldin-A (1:1000, Biolegend) and anti-CD107a–FITC (clone H4A3, BioLegend). Cells were incubated for 5 h, washed in cold PBS, and stained with live/dead Fixable Aqua (Thermo Fisher Scientific), anti-CD3–PECy7 (clone UCHT1, BioLegend) or anti-CD3-BV711 (Clone SK7, Biolegend), anti-CD56–BV605 (clone 5.1H11, BioLegend), anti-CD57–APC (clone HNK-1, BioLegend) or anti-CD57-PECy7 (clone HNK-1, biolegend), and anti-NKG2C–PE (clone 134591, R&D Systems). In some experiments, cells were also fixed/permeabilized using Cytofix/Cytoperm (BD Biosciences) and stained with anti-TNFα–BV421 (clone MAb11, BioLegend) and anti-IFNγ–APC (clone 4S.B3, BioLegend). Data were acquired using an AttuneNxT (Thermo Fisher) and analyzed with Attune NxT software or FlowJo software version 10 (Tree Star). Individual assays were run in technical triplicate. To ensure inter-assay variation did not affect results, a donor serum demonstrating moderate ADCC activity against SARS-CoV2 was included as a positive standard in every assay, while a serum collected before 2020 was included as a negative control in every assay. Where sera were tested at a range of dilutions, the area under the curve (AUC) was calculated using Graphpad Prism 9. This value was then normalised to the AUC for the standard serum in that particular assay. In all experiments, serums were additionally tested against mock infected cells to control for non-specific activation of NK cells by serum components. An example gating strategy is shown in figure S8.

### Serums

Serums were collected from healthy donors vaccinated with the BNT162b2 vaccine, a minimum of 3 weeks after the first dose, or 1 week after the second. A pre-vaccine sample was taken from every donor, and an ELISA for SARS-CoV2 RBD performed as previously described^88^, to determine pre-exposure to live SARS-CoV2. Longitudinal serums from naturally infected individuals have been described previously, clinical characteristics are in Table S2^50^. Those labelled ‘mild’ had severity scores of 0, 1, 2 or 3, and ‘severe’ was 4 or 5.

### Virus neutralisation assays

600PFU of SARS-CoV2 was incubated with appropriate dilutions of serum, in duplicate, for 1h, at 37°C. The mixes were then added to pre-plated VeroE6 cells for 48h. After this time, monolayers were fixed with 4% PFA, permeabilised for 15 min with 0.5% NP-40, then blocked for 1h in PBS containing 0.1% Tween (PBST) and 3% non-fat milk. Primary antibody (anti-nucleocapsid 1C7, Stratech, 1:500 dilution) was added in PBST containing 1% non-fast milk and incubated for 1h at room temperature. After washing in PBST, secondary antibody (anti-mouse IgG-HRP, Pierce, 1:3,000 dilution) was added in PBST containing 1% non-fat milk and incubated for 1h. Monolayers were washed again, developed using Sigmafast OPD according to manufacturers’ instructions, and read on a Clariostar Omega plate reader. Wells containing no virus, virus but no antibody, and a standardised serum displaying moderate activity were included as controls in every experiment. NT50 were calculated in Graphpad Prism 9.

### Antibody depletions

Anti-spike antibody was depleted from sera using magnetic bead conjugated spike trimer protein (Acrobiosystems). Beads were resuspended in PBS+0.05% BSA, then 50µl serum was mixed with 150µl beads. Mixtures were incubated on a rotating mixer at 4°C overnight. Serum diluted in buffer alone was used as a control. Magnetic beads were then removed using a 3D printed magnetic stand. All values given in assays are corrected for this initial 4-fold dilution. Levels of anti-spike antibody were measured using a spike trimer ELISA (Acrobiosystems).

### Ethics

PBMC were extracted from apheresis cones obtained from the Welsh Blood Service (WBS) via an ad-hoc agreement or from blood samples from healthy volunteers and stored in liquid N_2_ until use. Use of healthy volunteer PBMC for this project, including those from WBS, was ethically approved by the Cardiff University School of Medicine Research Ethics Committee (SMREC) nos. 20/55 and 20/101. Recruitment of healthy volunteers after vaccination was covered by the Cardiff University School of Medicine Research Ethics Committee under reference no. 18/04.

---

**Figure S1.**
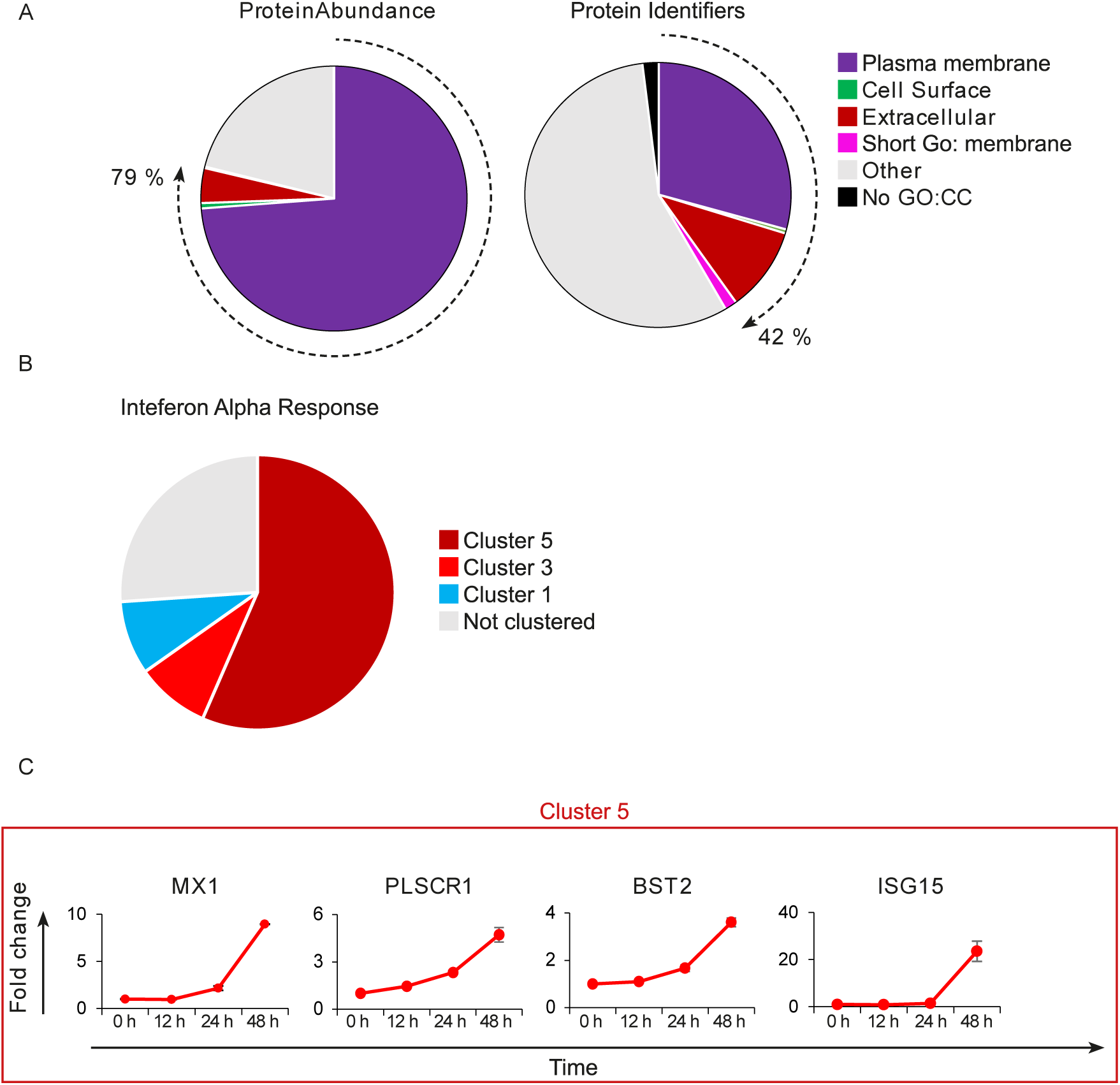
Plasma Membrane Proteins are enriched in the PMP dataset. (A) Pie charts showing the proportion of proteins falling into previously defined annotation categories for plasma membrane proteins by left, protein abundance, or right, protein identifiers. (B) Pie chart showing the distribution of proteins defined by the Molecular Signatures Database (MSigDB) gene set “Hallmark Interferon Alpha Response” into the defined temporal clusters. C. Example temporal profiles of type I interferon inducible genes upregulated in cluster 5, y-axis shows fold change compared to 0 h timepoint.

**Figure S2.**
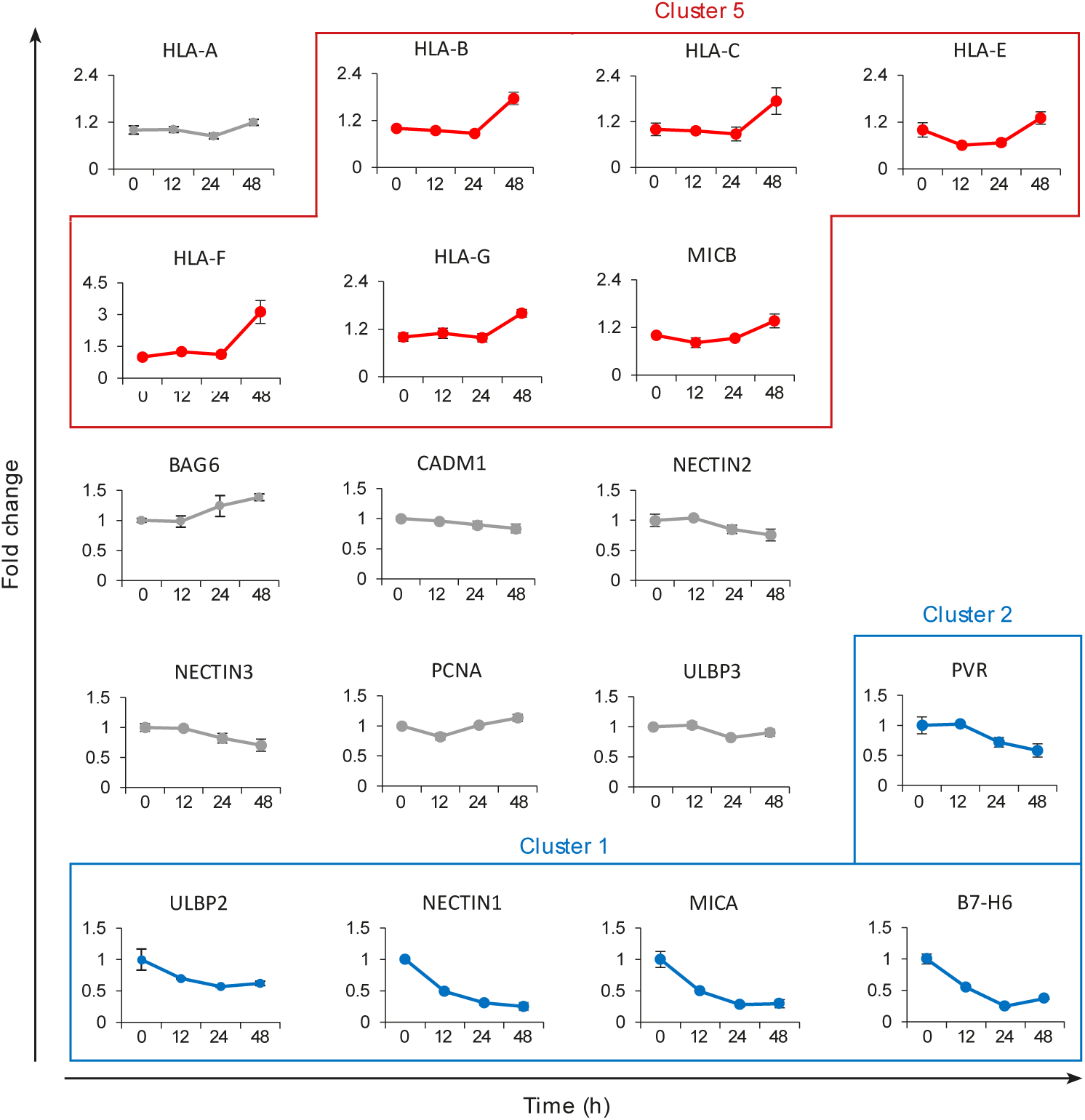
NK ligands detected in the PMP Dataset. Temporal profiles of NK-cell ligands detected in the PMP dataset, y-axis shows fold change compared to 0 h timepoint.

**Figure S3.**
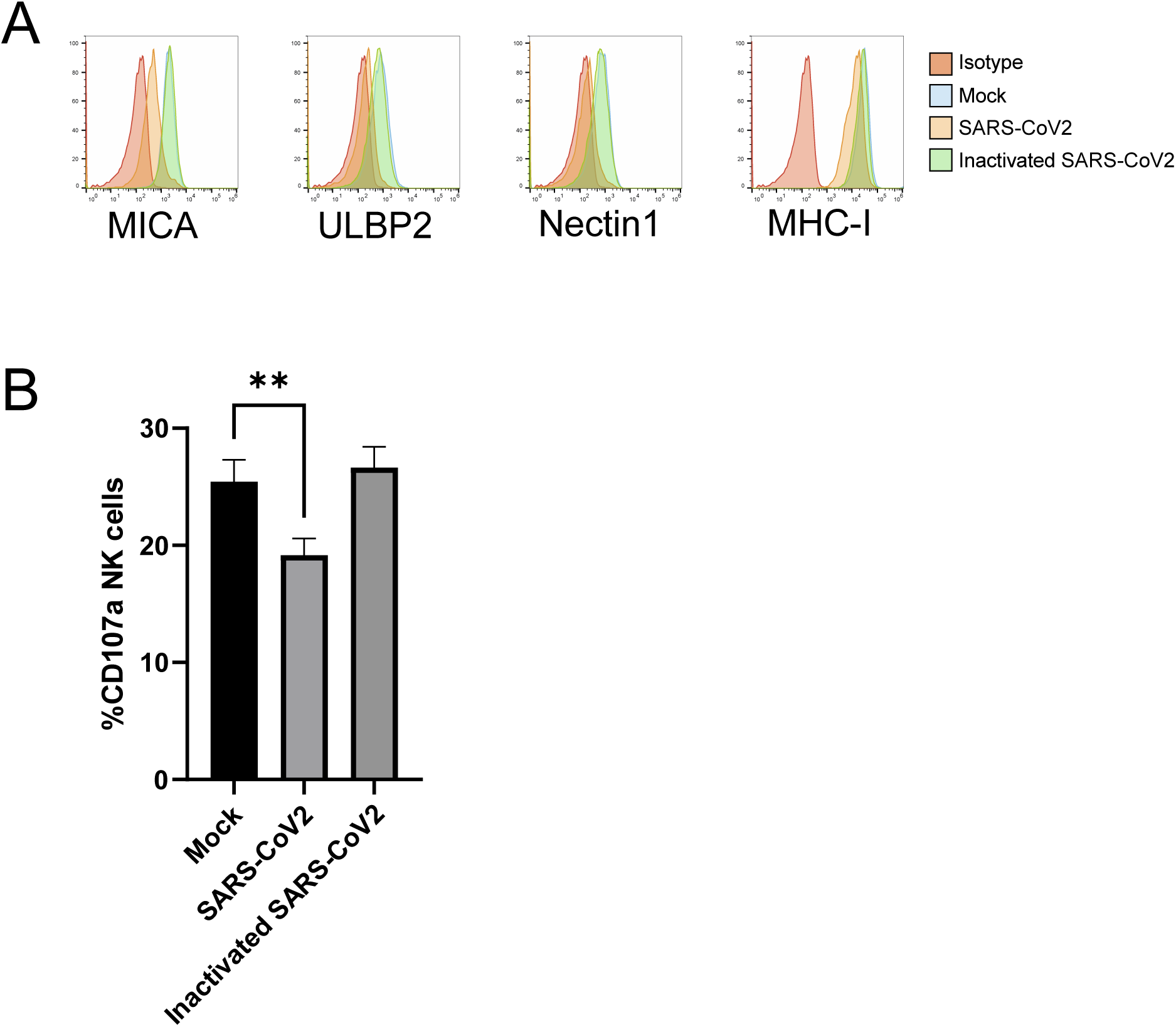
Modulation of NK activity is dependent on virus replication. AAT cells were mock infected, or infected with SARS-CoV2 (MOI=5), or with heat inactivated SARS-CoV2. After 24h, cells were dissociated with trypLE. (A) cells were stained with the indicated antibodies before being analysed by flow cytometry. (B) Cells were mixed with PBMC and incubated for 5h in the presence of golgistop and CD107a antibody, before staining for CD3, CD56, CD57 and Live/Dead Aqua. Cells were gated on live CD57+ NK cells, and the percentage of cells positive for CD107a calculated. Assays were run in technical triplicate. One-way ANOVA **p<0.01.

**Figure S4.**
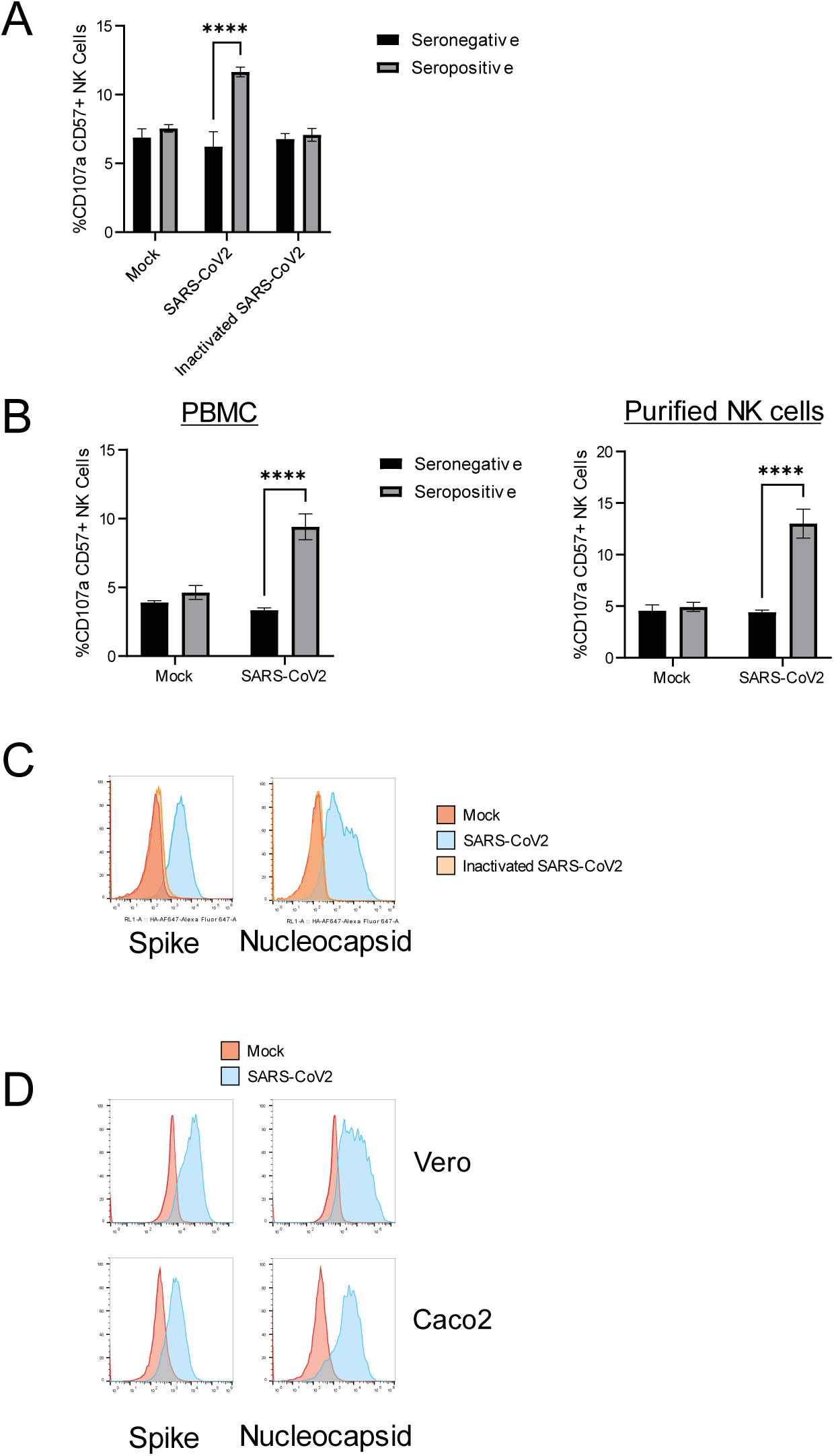
ADNKA is dependent on virus replication, and occurs through direct action on NK cells. (A-C) AAT cells were mock infected, or infected with SARS-CoV2, or with heat inactivated SARS-CoV2 (MOI=5). After 24h, cells were dissociated with trypLE. (A-B) Cells were mixed with PBMC (A) or purified NK cells (A-B) and 1% sera that was seropositive or seronegative for SARS-CoV2. Cells were incubated for 5h in the presence of golgistop and CD107a antibody, before staining for CD3, CD56, CD57 and Live/Dead Aqua. Cells were gated on live CD57+ NK cells, and the percentage of cells positive for CD107a. Assays were run in technical triplicate. Two-way ANOVA ****p<0.0001. (C) cells were stained for Spike or Nucleocapsid. (D) Vero or Caco2 cells were infected with SARS-CoV2. After 24h, cells were dissociated with trypLE and stained for Spike or Nucleocapsid, before being analysed by flow cytometry.

**Figure S5.**
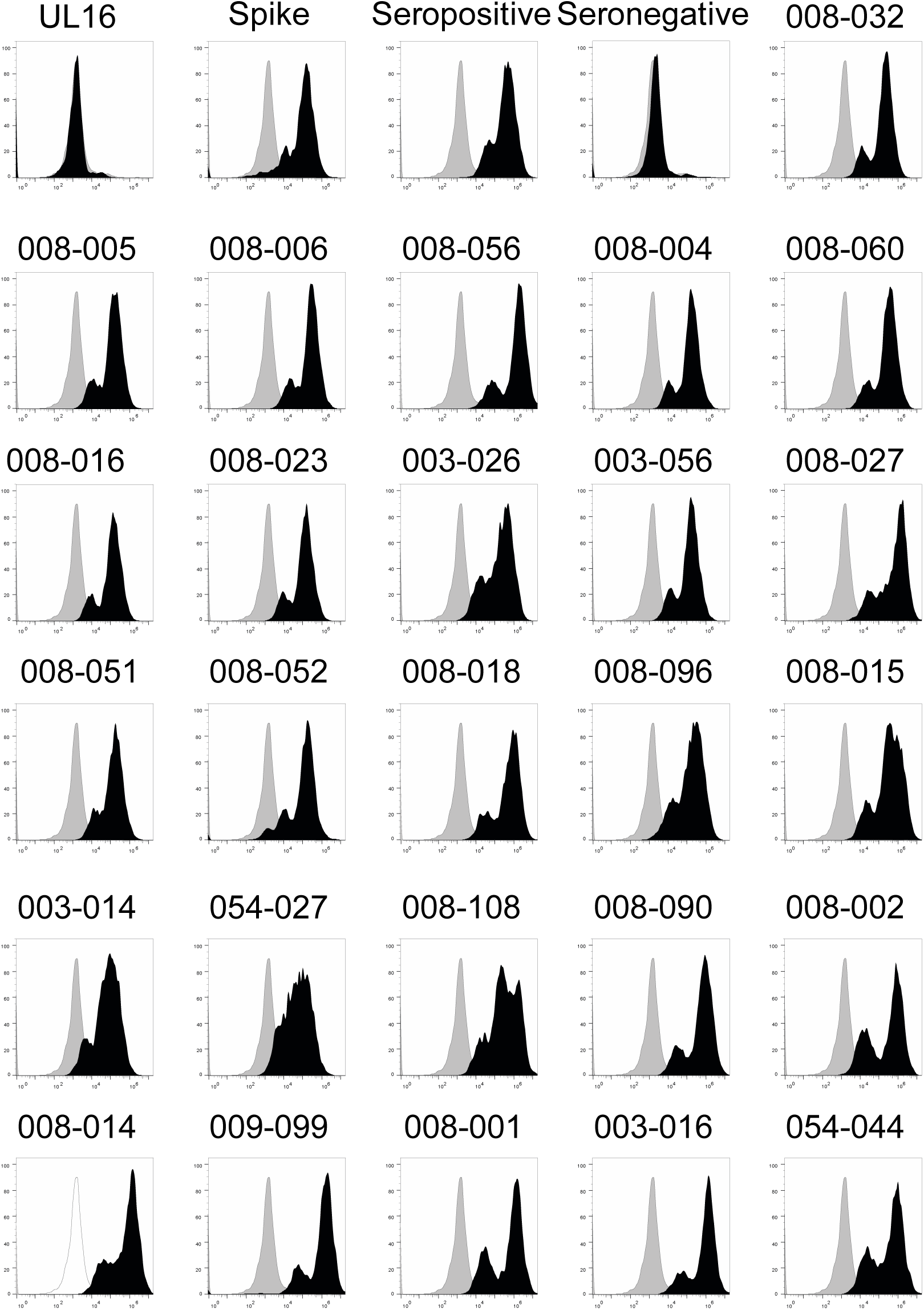
Anti-spike antibodies bind to SARS-CoV2 infected cells. AAT cells were mock infected (light grey), or infected with SARS-CoV2 (black; MOI=5) for 24h, then detached with TrypLE, stained with the indicated human anti-spike antibodies, then analysed by flow cytometry.

**Figure S6.**
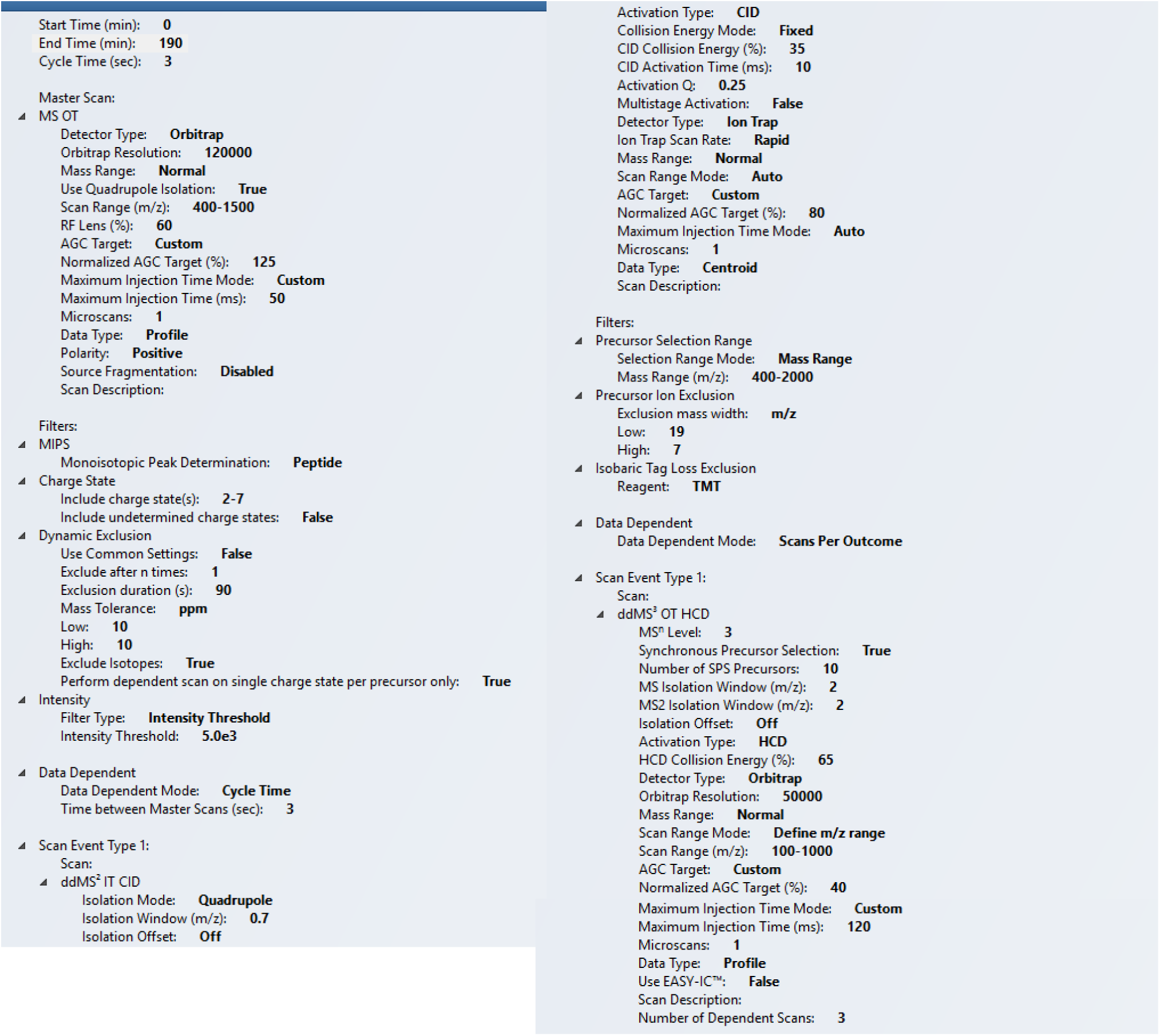
MS Method parameters.

**Figure S7.**
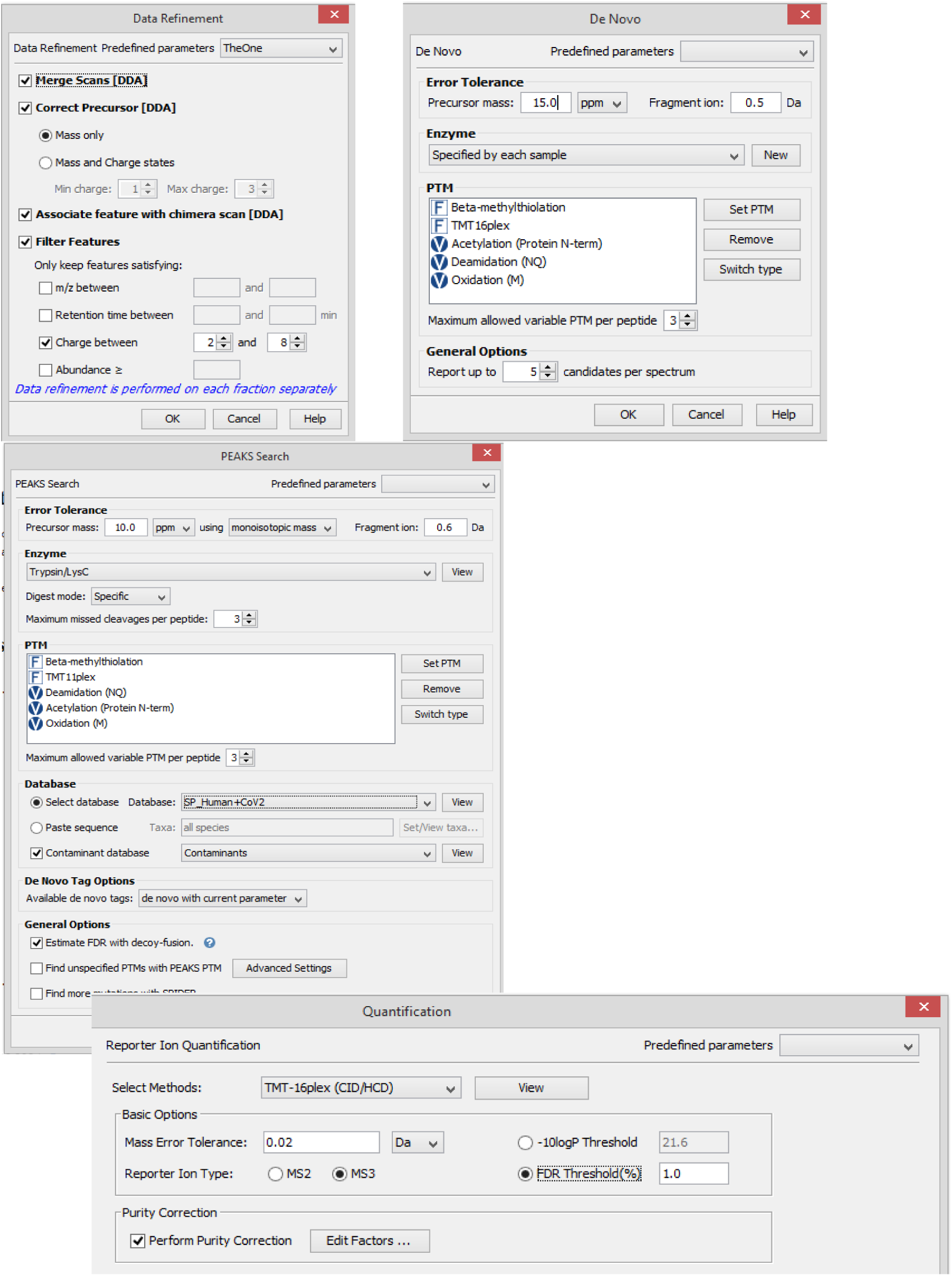
Search Parameters for MS data processing.

**Figure S8.**
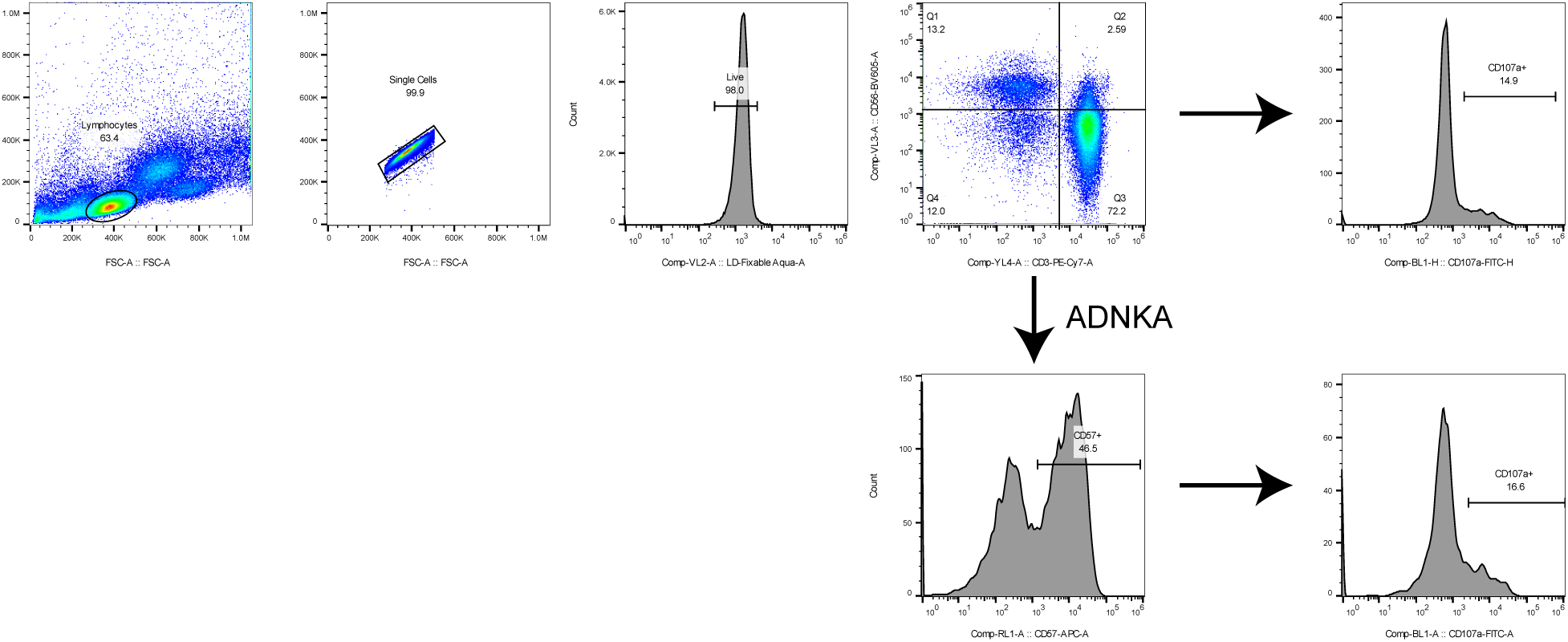
Example gating strategy for NK assays. Lymphocytes were gated on FSC/SSC, doublets excluded using FSC-A/FSC-H, and live cells selected on the basis of Live/Dead Aqua staining. The percentage of CD107a+ cells were determined in the CD56+/CD3- gate, or in the CD57+ gate of CD56+/CD3- cells.

**Table S2.** Clinical characteristics of patients giving longitudinal serum samples

| Study ID | Severity | Age | Gender | Past medical history |
| --- | --- | --- | --- | --- |
| 1 | Mild | 82 | M | dementia, ESRF, HTN, IHD, PVD |
| 2 | Mild | 77 | F | ESRF, HTN, T2DM |
| 3 | Mild | 64 | F | AF, CVA, dementia, ESRF, HTN, T2DM |
| 4 | Mild | 27 | F | ESRF, obesity |
| 5 | Mild | 48 | F | ESRF, HTN, obesity |
| 6 | Severe | 52 | F | Asthma, obesity |
| 7 | Severe | 62 | M | colon ca, epilepsy, HTN, obesity, T2DM |
| 8 | Severe | 65 | F | HTN, hypothyroidism, obesity |
| 9 | Severe | 61 | M | ESRF, obesity, VTE |
| 10 | Severe | 57 | M |  |
| 11 | Mild | 23 | F | CKD, HTN, T1DM |
| 12 | Mild | 62 | M | ESRF, HTN, renal transplant |
| 13 | Mild | 25 | F | AML |
| 14 | Mild | 52 | M | CVA, prostate ca, T2DM |
| 15 | Mild | 88 | F | AF, CCF, CKD, COPD, IHD, RA, T2DM |
| 16 | Mild | 64 | M |  |
| 17 | Mild | 69 | M | AF, HTN, T2DM, VTE |
| 18 | Mild | 22 | M | NHL |
| 19 | Severe | 49 | M |  |
| 20 | Severe | 67 | F | AF, HTN, obesity, T2DM |

AF=atrial fibrillation, AML=acute myeloid leukaemia, ca=cancer, CCF=congestive cardiac failure, CKD=chronic kidney disease, COPD=chronic obstructive pulmonary disease, CVA cerebrovascular accident, ESRF=end-stage renal failure, HTN=hypertension, IHD=ischemic heart disease, NHL=non-Hodgkin lymphoma, RA=rheumatoid arthritis, T1DM/T2DM=type 1/2 diabetes mellitus, VTE=venous thromboembolism.

**Table S3.** Monoclonal Ant-spike Antibodies used, data taken from^50^

| mAb name | Specificity | Competition group | S EC50 | RBD EC50 | NTD EC50 | S2 ELISA | SARS-CoV-2 WT FL IC50 | SARS-CoV-2 WT PV IC50 |
| --- | --- | --- | --- | --- | --- | --- | --- | --- |
| P008_060 | NON-S1 | 7 | 0.031 | >10 | >10 | n.d. | 32.90 | 0.095 |
| P008_004 | NON-S1 | n.d. | 0.045 | >10 | >10 | + | >100 | >100 |
| P008_005 | NON-S1 | n.d. | 0.20 | >10 | >10 | + | >100 | >100 |
| P008_006 | NON-S1 | n.d. | 0.10 | >10 | >10 | + | >100 | >100 |
| P008_016 | NON-S1 | n.d. | 0.11 | >10 | >10 | + | >100 | >100 |
| P008_023 | NON-S1 | n.d. | 0.40 | >10 | >10 | + | >100 | >100 |
| P008_032 | NON-S1 | n.d. | 0.060 | >10 | >10 | + | >100 | >100 |
| P008_051 | NTD | 5 | 0.091 | >10 | 0.056 | n.d. | 48.82 | 0.30 |
| P008_052 | NTD | 5 | 0.085 | >10 | 0.063 | n.d. | 48.65 | 0.13 |
| P003_027 | NTD | 6 | 0.0039 | >10 | 0.0065 | n.d. | 1.64 | 12.92 |
| P008_056 | NTD | 6 | 0.060 | >10 | 0.049 | n.d. | 0.014 | >100 |
| P008_001 | NTD | n.d. | 0.82 | >20 | 1.20 | n.d. | >100 | >100 |
| P008_002 | NTD | n.d. | 0.0084 | >20 | 0.015 | n.d. | >100 | >100 |
| P008_014 | NTD | n.d. | 0.066 | >20 | 0.021 | n.d. | 1.81 | >100 |
| P008_099 | NTD | n.d. | 0.022 | >20 | 0.15 | n.d. | >100 | >100 |
| P054_044 | NTD | n.d. | 0.031 | >20 | 4.00 | n.d. | 0.94 | 0.74 |
| P003_016 | NTD | n.d. | 0.13 | >20 | 2.00 | n.d. | 0.15 | 0.45 |
| P003_014 | RBD | 1 | 0.0076 | 0.0086 | n.d. | n.d. | >100 | >100 |
| P054_027 | RBD | 1 | 0.0077 | 0.013 | n.d. | n.d. | >100 | 0.7 |
| P008_018 | RBD | 3 | 0.071 | 0.082 | n.d. | n.d. | 0.055 | 0.00081 |
| P008_090 | RBD | 3 | 0.018 | 0.026 | n.d. | n.d. | 0.071 | 0.0042 |
| P008_108 | RBD | 3 | 0.0092 | 0.014 | n.d. | n.d. | 0.0023 | 0.0031 |
| P008_015 | RBD | 4 | 0.034 | 0.036 | n.d. | n.d. | 0.055 | 0.00012 |
| P008_096 | RBD | 4 | 0.030 | 0.055 | n.d. | n.d. | 9.09 | 0.41 |

